# A family of E3 ligases extend K11 polyubiquitin on sites of MARUbylation

**DOI:** 10.1101/2025.05.11.653360

**Authors:** Rachel E Lacoursiere, Kapil Upadhyaya, Jasleen Kaur Sidhu, Daniel S Bejan, Ivan Rodriguez Siordia, Michael S Cohen, Jonathan N Pruneda

## Abstract

Ubiquitin (Ub) cooperation with other post-translational modifications provides a tiered opportunity for protein regulation. Small modifications to Ub such as phosphorylation, acetylation, or ADP-ribosylation have varying impacts on signaling. The Deltex family of E3 ligases was previously implicated in the ubiquitylation of ADP-ribose (ADPr) and ADPr-containing macromolecules. Our previous work found ester-linked mono-ADPr ubiquitylation (MARUbylation) on PARP7 and PARP10 in cells and that this mark is extended with K11 polyUb. We previously screened for E3 ligases that interact with PARP7 through three different approaches and identified six candidates, including the Deltex family member DTX2. One of these hits, RNF114, interacts with various other PARPs, leading us to hypothesize that RNF114 binds to sites of MARUbylation and extends K11 polyUb. Here, we show that DTX2 generates the initial MARUbe on PARP7 in cells, which depends on PARP7 catalytic activity. The MARUbe on PARP7 is extended with K11 polyUb by RNF114. To investigate the mechanism of RNF114 reader/writer function, we developed a click chemistry-inspired chemoenzymatic approach to create a novel fluorescent Ub-ADPr probe for studying its interaction with RNF114. Strikingly, we found that RNF114 has a weak affinity for ADPr and Ub separately but explicitly recognizes the linkage between Ub and ADPr present in MARUbylated species. We used AlphaFold3 modeling to examine the mechanisms of Ub-ADPr recognition and K11-linked polyUb extension by RNF114. We identified a tandem Di19-UIM module in RNF114 as a <u>M</u>AR<u>U</u>be-<u>b</u>inding <u>d</u>omain (MUBD), thus providing a reader function that interfaces with K11-specific writer activity. Finally, we described a small family of MUBD-containing E3 ligases that demonstrate preference for Ub-ADPr, which we call <u>M</u>ARUbe-<u>T</u>argeted <u>L</u>igases (MUTLs).

## Introduction

Ubiquitylation is one of the most prevalent and versatile post-translational modifications (PTMs) in the eukaryotic cell. Concerted efforts from three classes of ubiquitylating enzymes (E1 activating enzymes, E2 conjugating enzymes, and E3 ligases) facilitate the covalent modification of a substrate with the small protein ubiquitin (Ub). Canonically, ubiquitylation occurs at lysine sidechains in a proteinaceous substrate, and repeated cycles of the cascade can result in the formation of polyubiquitin (polyUb) chains through Ub-Ub linkages (Swatek & Komander, 2016).

As a protein itself, Ub is susceptible to regulation by other PTMs such as phosphorylation and acetylation. For example, phosphorylation of Ub at Ser65 by the kinase PINK1 acts as a crucial initial signal for Parkin/PINK1-dependent mitophagy (Kane et al., 2014; Kazlauskaite et al., 2014; Koyano et al., 2014). Acetylation of Ub lysine residues alters the efficiency of E2∼Ub conjugate formation (Lacoursiere & Shaw, 2021). Further, Ub signaling can be modulated through the activity of bacterial effectors that target Ub for ADP-ribosylation at various sidechains (Qiu et al., 2016; Yan et al., 2020). Human enzymes including PARPs and the Deltex family of E3 ligases have also been reported to ADP-ribosylate Ub, though this modification occurs at the C-terminus (Ashok et al., 2022; Chatrin et al., 2020; Yang et al., 2017). While the exact role of this modification is unclear, interplay between ADP-ribosylation and ubiquitylation is prevalent in pathways such as the DNA damage response (DDR) and innate immune signaling (Pellegrino & Altmeyer, 2016).

Crosstalk between ADP-ribosylation and ubiquitylation can be exemplified by the poly-ADP-ribosylation (PARylation)-dependent ubiquitylation (PARdU). RNF146, which complexes with the PARylating PARPs Tankyrase1/2 (PARP5a/b), facilitates the canonical ubiquitylation of a substrate in a PARylation-dependent manner (Callow et al., 2011; Zhang et al., 2011). PAR recognition occurs at the internal repeating unit *iso*-ADPr by the Trp-Trp-Glu (WWE) PAR “reader” domain of RNF146, which induces a conformational change and places the catalytic E3 ligase RING domain in an active orientation to facilitate catalysis (DaRosa et al., 2015). PARdU demonstrates the cooperation between PARPs and E3 ligases in an ADP-ribosylation dependent manner. Beyond RNF146, other E3 ligases contain WWE domains, including the Deltex RING E3 ligases DTX1, DTX2, and DTX4 (Chatrin et al., 2020). Deltex E3 ligases are also characterized by a Deltex carboxy-terminal (DTC) domain at their C-termini. Through *in vitro* experiments with select family members, this domain has been shown to bind and ubiquitylate NAD^+^, free ADPr, or even mono-ADP-ribosylated (MARylated) substrates such as proteins and nucleic acids (Ahmed et al., 2020; Chatrin et al., 2020; Zhu et al., 2024; Zhu et al., 2022).

We recently discovered that ubiquitylation of MAR protein modifications occurs in cells as a dual PTM downstream of multiple PARPs under conditions including interferon stimulation (Bejan et al., 2025). We termed this process MAR ubiquitylation (MARUbylation) to distinguish it from MARylation and canonical ubiquitylation. The first step in MARUbylation requires PARP catalytic activity to attach mono-ADP-ribose to a protein substrate, like itself. Next, an E3 Ub ligase recognizing ADPr and/or the MARylated substrate ubiquitylates the MAR through a labile ester linkage, forming the MAR-Ub ester (MARUbe). Although Deltex E3 ligases have not been formally shown to generate the initial MARUbe in cells, an abundance of *in vitro* experiments supports this hypothesis. Interestingly, we identified K11 polyUb linkages extending from the initial MARUbe, suggesting the activity of an additional K11-specific ligase. Together, MARUbylation represents three layers of PTM regulated by at least as many enzymes.

Multiple lines of evidence connect PARPs with Deltex E3 ligases, providing confidence in a functional role for MARUbylation in cells (Huttlin et al., 2021; Oughtred et al., 2021; Schweppe et al., 2018). The same lines of evidence also identify RNF114 and/or RNF166, two RING-type E3 ligases with related sequence and domain topologies. Previous work has shown that RNF114 responds to ADP-ribose signaling. For example, RNF114 is recruited to sites of DNA damage in an ADP-ribosylation-dependent manner (Djerir et al., 2024; Li et al., 2023). Furthermore, outside of the RING domain, both RNF114 and RNF166 encode Zn^2+^-coordinating drought-induced 19 (Di19) domains that have been shown to interact with ADPr (Longarini et al., 2023), and Ub-interacting motifs that recognize Ub (Giannini et al., 2008). Considering this, we hypothesized that RNF114 may read MARUbe modifications downstream of PARP and Deltex activity using its tandem Di19-UIM module, and using the catalytic RING domain, append K11 polyUb as a third layer of PTM regulation. Indeed, in cellular knockdown experiments, we find that PARP, Deltex, and RNF114 activity are all required for K11-extended MARUbylation. Using a newly generated fluorescent Ub-ADPr substrate, we validated specific recognition and K11 polyUb extension by RNF114. To reveal the mechanism of RNF114 reader/writer function, we used AlphaFold3 to model multi-protein complexes and rationalized RNF114 mutations that validate these functions. Lastly, we found that RNF114 and RNF166 are part of a small family of tandem Di19-UIM module-containing E3 ligases. We term this module the <u>M</u>AR<u>U</u>be-<u>b</u>inding <u>d</u>omain (MUBD) and show that E3 ligases possessing this domain can act as <u>M</u>AR<u>U</u>be-targeted ligases (MUTLs).

## Results

### DTX2 and RNF114 regulate the formation and extension of MARUbe on PARP7 in cells

Our recent studies revealed the presence of a unique dual PTM, MARUbe, on PARP7 and PARP10 in cells (Bejan et al., 2025). We also found that MARUbe was extended with K11-linked polyUb. We first sought to determine the identity of the amino acids conjugated to MARUbe on PARP7. In the case of PARP10, MARUbe is attached to Glu/Asp amino acids since NH_2_OH and the Glu/Asp-specific MARylases, MacroD1 and MacroD2, release a K11-linked polyUb MARUbe species from immunoprecipitated GFP-PARP10 (Bejan et al., 2025). When it comes to PARP7, it is a bit more complicated because PARP7 transfers ADPr from NAD^+^ to multiple amino acids, including Glu/Asp and Cys (Gomez et al., 2018; Palavalli Parsons et al., 2021; Rodriguez et al., 2021). It is important to note that the relative abundance of Glu/Asp MARylation versus Cys MARylation is difficult to address due to the lability of Glu/Asp MARylation (Tashiro et al., 2023; Weixler et al., 2023). Identifying which amino acids contain the MARUbe mark on PARP7 is an essential first step to understand its regulation in cells.

To determine if Glu/Asp or Cys are the predominant sites of PARP7 MARUbylation, we used a mutagenesis approach. We expressed WT GFP-PARP7 as well as a mutant in which five of the most abundant Cys-auto-MARylation sites−identified by MARylation amino acid site proteomics (Rodriguez et al., 2021)−were mutated to Ser (C5: C39, C294, C351, C543, C552) in PARP1 knockout (KO) HEK 293 cells. PARP1 KO cells were used to eliminate background ADP-ribosylation mediated by abundantly expressed PARP1. As expected based on our proteomics data, the C5 PARP7 mutant exhibited substantially lower auto-MARylation levels compared to WT PARP7 (Fig. 1A). This result confirms that these five Cys residues are the major Cys-auto-MARylation sites on PARP7.

**Figure 1.**
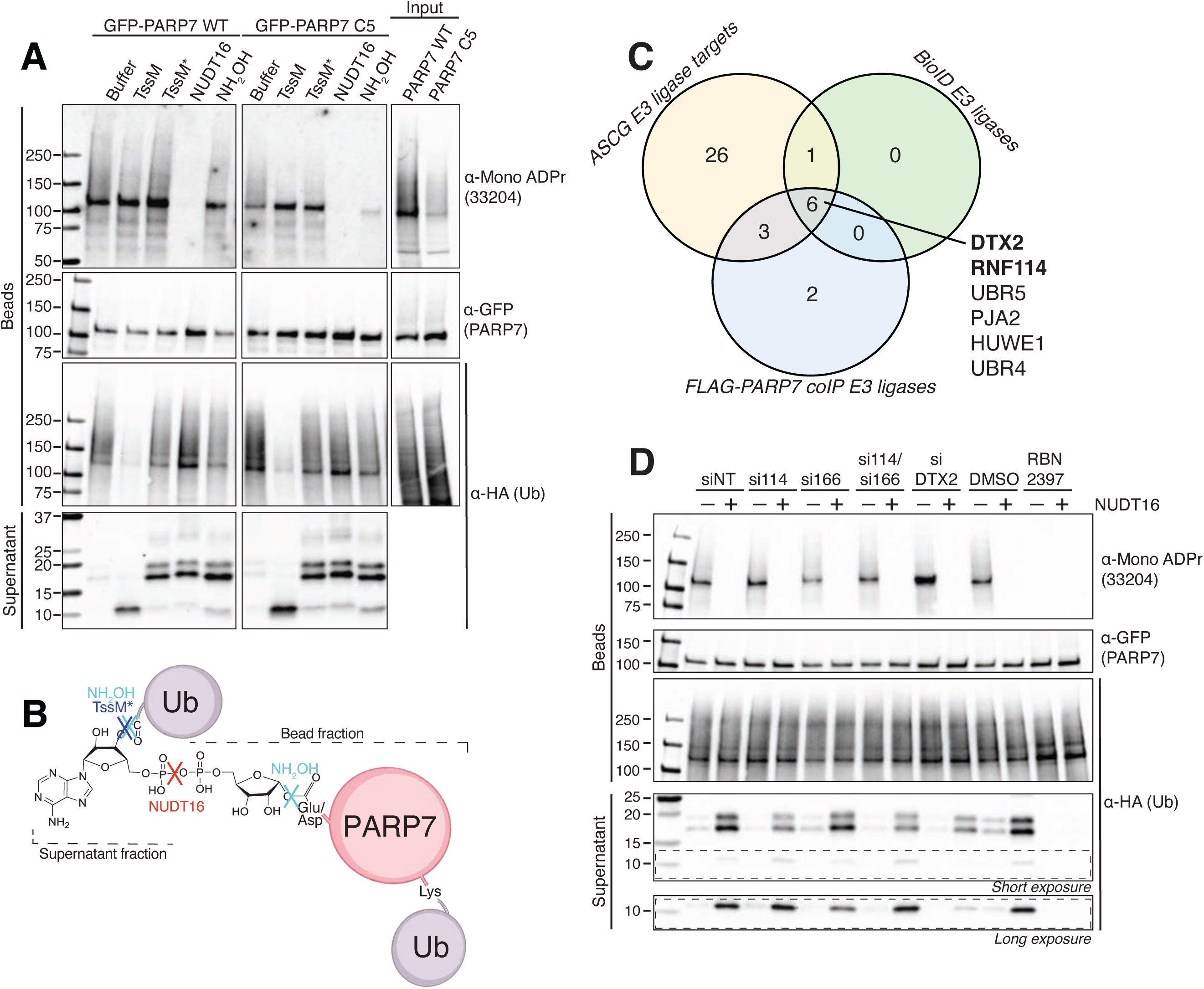
RNF114 extends PARP7 MARUbylation on Glu/Asp residues. **A**) A MARUbylation assay for GFP-PARP7 WT or the C5 mutant of PARP7, where all cysteines have been mutated to serines, performed in HEK 293 PARP1 KO cells. GFP-PARP7 pulldowns were subjected to treatment with TssM, TssM*, NUDT16, or hydroxylamine (NH_2_OH) treatment and evaluated for the composition of the bound vs. supernatant fractions. All samples from this experiment were present on the same membrane. PARP7 WT and C5 samples were imaged together but cropped for clarity to remove an intervening lane. The input samples were imaged at a shorter exposure to avoid signal saturation. **B**) Schematic representation of a MARUbylation assay using NUDT16, TssM* or NH_2_OH to cleave the MAR-bound Ub. **C**) Identification of six candidate E3 ligases common to three approaches to identify ligases that interact with PARP7. **D**) GFP-PARP7 C5 MARUbylated species were pulled down from HEK 293T cells following inhibitor or siRNA treatment as indicated and subjected to a MARUbylation assay using NUDT16 to separate the stable PARP7 canonical ubiquitylation from the labile MARUbylation. DMSO or NT (non-targeting siRNA) were used as controls for the RBN2397 PARP7 inhibitor treatment and siRNA knockdowns, respectively.

We then determined the relative levels of MARUbylation for WT and C5 PARP7 using our IP MARUbylation assay (Bejan et al., 2025). Briefly, the IP MARUbylation assay involves transiently expressing tagged Ub (HA-Ub) together with a GFP-tagged WT or C5 PARP7 in PARP1 KO HEK 293 cells. The GFP-tagged PARP7 proteins are immunoprecipitated using GFP-trap beads. Stringent washing of beads is performed under denaturing conditions (7M urea and 1% SDS) to remove noncovalently bound Ub. We then perform on-bead enzymatic (TssM*, a Ub esterase or NUDT16, an ADPr pyrophosphatase) or chemical treatment (NH_2_OH, cleaves Ub ester and Glu/Asp MARylation) to elute the K11-extended MARUbe species into the supernatant, which can be detected by western blotting by probing for the HA tag (Fig. 1B). With these treatments, canonical Lys-linked ubiquitylation remains attached to GFP-tagged PARP7 and can be detected in the bead fraction. Using this assay found that the K11-extended MARUbe levels are similar between WT and C5 PARP7 (Fig. 1A). These results show that Cys residues are not the major sites of PARP7 MARUbylation. We speculate that Glu/Asp residues are the major sites of PARP7 MARUbylation.

After identifying the specific amino acids MARUbylated on PARP7, identifying the E3 ligases responsible for the formation and K11-extension of the MARUbe modification remained. Addressing this knowledge gap is critical for elucidating the cellular function of MARUbylation. We therefore took a candidate approach, focusing on PARP7 MARUbylation because of the good availability of data on interactors and targets. Moreover, PARP7 is a highly sought-after therapeutic target in cancer (Gozgit et al., 2021) and stroke (Cai et al., 2024) and deciphering the function of PARP7-mediated MARUbylation is critical for understanding its cellular function in these diseases. To identify potential E3 ligases that regulate MARUbylation, we analyzed mass spectrometry (MS) data from three sources: 1. BioID proximity labeling (Rodriguez et al., 2021), 2. Co-IP using FLAG-PARP7 (Zhang et al., 2020), and 3. An analog sensitive chemical genetics (ASCG) approach for identifying direct PARP7 MARylation targets (Data unpublished). Six E3 ligases were identified in all three MS datasets (Fig. 1C). DTX2 contains a MAR-binding domain (MBD), and was shown by us and others to attach Ub to the 3’OH of the adenosine ribose on a wide array of ADPr-containing species *in vitro* (Ahmed et al., 2020; Bejan et al., 2025; Dearlove et al., 2024; Kelly et al., 2024; Zhu et al., 2024; Zhu et al., 2022). Likewise, recent studies have shown that RNF114 contains a MBD (Longarini et al., 2023), and BioGRID 4.4 and BioPlex interactome analyses showed that RNF114, and the closely related RNF166, are top interactors of several MARylating PARPs, namely PARP7, PARP10, PARP11, and PARP14 (Huttlin et al., 2021; Oughtred et al., 2021; Schweppe et al., 2018). We therefore hypothesized that DTX2 catalyzes the initial MARUbe mark, and that RNF114 and/or RNF166 were responsible for the subsequent extension of K11 polyUb.

To determine if DTX2 and RNF114/RNF166 are involved in the formation and extension of MARUbe on PARP7 in PARP1 KO HEK 293 cells, we used an siRNA-mediated approach and our IP MARUbylation assay with C5 GFP-PARP7. We first confirmed that C5 PARP7 MARUbylation was indeed dependent on PARP7 activity by treating transfected cells with the PARP7-specific inhibitor RBN2397 (Fig. 1D, Supplemental Fig. 1) (Gozgit et al., 2021). RBN2397 eliminated MARylation as well as the K11-extended MARUbe species, demonstrating that C5 PARP7 MARUbylation is dependent on PARP7 catalytic activity. Knockdown of DTX2 reduced both the MARUbe itself (∼10 kDa) and the K11-extended MARUbe species (15-20 kDa), indicating that DTX2 regulates the initial ubiquitylation of MAR on PARP7 (Fig. 1D). Upon siRNA knockdown of RNF114, we observed a reduction in the K11-extended MARUbe species, but an increase in MARUbe (Fig. 1D). Surprisingly, we did not observe this effect upon knockdown of RNF166 (Fig. 1D). Together these results support our hypothesis that DTX2 is the major PARP7 MARUbe-generating E3 ligase and RNF114 is the major E3 ligase that extends MARUbe with K11 polyUb.

### An integrated click chemistry and enzymatic strategy yields a novel fluorescent Ub-ADPr tool

Having established RNF114 as a regulator of PARP7 MARUbylation in cells, we next aimed to characterize its *in vitro* activity to understand how RNF114 catalyzes K11-linked ubiquitylation of MARUbe. We envisioned that an ester-linked, fluorescent Ub-ADPr conjugate could serve as a valuable probe for fluorescence polarization (FP)-based assays, enabling real-time monitoring of RNF114 activity, similar to Ub-Lys-TAMRA (Ub-Lys^T^) for studying canonical ubiquitylation (Geurink et al., 2012). We developed a chemoenzymatic approach to produce a fluorescent ester-linked Ub-ADPr conjugate (Fig. 2A). We first synthesized a clickable ADPr analog containing an α*-O*-propargyl at the anomeric carbon of the ribose ring (compound **8,** OP-ADPr) as described previously, but with minor modifications (Supporting Information) (Liu et al., 2018). We next coupled Ub to OP-ADPr using DTX2 to generate an ester-linked Ub-OP-ADPr (Materials and Methods). Ub-OP-ADPr was purified using ion exchange chromatography while monitoring the absorbance at 280 nm (protein) and 254 nm (ADPr). Consistent among many preparations, Ub-OP-ADPr eluted separately from excess Ub and other reaction components on a Resource S column (Fig. 2A, chromatogram inset). Samples of Ub before and after conjugation to OP-ADPr-were evaluated by intact mass spectrometry (Fig. 2B). The observed mass for Ub (8564.4 Da) was in good agreement with the theoretical mass (8564.8 Da). After conjugation to OP-ADPr, the observed mass was 9143.3 Da, in agreement with the expected mass of 9126.9 Da. We attribute the extra ∼16 Da of mass to methionine oxidation of Ub over time. To generate the fluorescent Ub-ADPr analog (Ub-ADPr^T^), Ub-OP-ADPr was coupled to TAMRA-azide using Copper-Catalyzed Azide-Alkyne Cycloaddition (CuAAC, “click chemistry”). The Ub-ADPr^T^ species was resolved from any unreacted TAMRA-azide using size exclusion chromatography, and SDS-PAGE confirmed generation of the Ub-ADPr^T^ with in-gel fluorescence (Fig. 2C).

**Figure 2.**
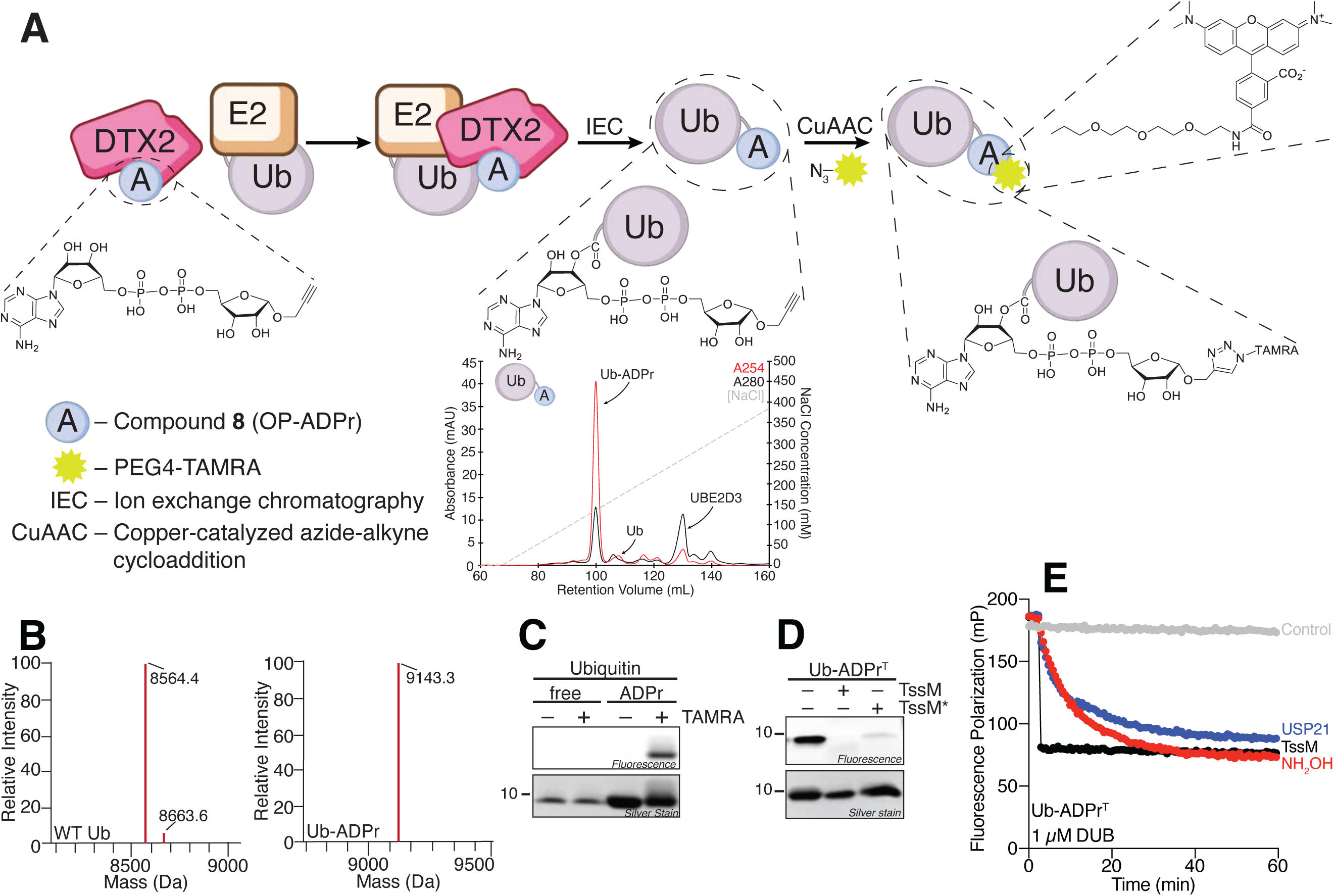
Synthesis of fluorescent Ub-ADPr (Ub-ADPr^T^) and ADPr^T^ species. **A**) Schematic representation of the workflow used to produce the fluorescent molecule Ub-ADPr^T^. The chromatogram provides a representative example for the isolation of Ub-ADPr from other reaction components by ion exchange chromatography. **B**) Validation of Ub-ADPr by intact mass spectrometry. WT Ub MW_expected_= 8564.84 Da (top), Ub-ADPr MW_expected_= 9126.9 Da (bottom). **C**) Validation of Ub-ADPr^T^ using in-gel fluorescence before and after the click reaction. **D**) In-gel fluorescence analysis of a separate experiment treating Ub-ADPr^T^ with TssM or the Ub esterase TssM*. **E**) Validation of Ub-ADPr^T^ through fluorescence polarization experiments with the indicated deubiquitylase treatments.

The final Ub-ADPr^T^ product was validated by SDS-PAGE after treatment with TssM or the Ub esterase TssM*, which confirmed the expected loss in fluorescence (Fig. 2D). We further validated Ub-ADPr^T^ in a fluorescence polarization experiment using hydroxylamine and the nonspecific deubiquitylases TssM and USP21 to confirm that 1) the Ub is attached to ADPr via an ester linkage; 2) there were no nonspecific interactions between the Ub and TAMRA-azide, and that the fluorophore was on the ADPr-side of the MARUbe; and 3) that fluorescence polarization was a valid approach to monitor the state of the Ub-ADPr^T^ substrate (Fig. 2E). Together, these experiments not only validate the identity of Ub-ADPr^T^, but also that Ub-ADPr^T^ is a useful FP probe.

In parallel, we chemically synthesized ADPr^T^ as described in the Supporting Information and validated it against the known ADPr-binding protein AF1521 (Karras et al., 2005). Using the ADPr^T^ substrate in a fluorescence polarization assay, we determined the K_d_ of its interaction with AF1521 to be 1.2 µM (Supplemental Fig. 2), which is in good agreement with the previously reported K_d_ of 3 µM (Nowak et al., 2020). These findings confirm that the TAMRA fluorophore does not interfere with the activity of deubiquitylases or canonical ADPr-binding proteins, so its positioning in Ub-ADPr^T^ would not be expected to interfere with other protein-protein interactions.

### RNF114 binds ADPr and Ub

RNF114 encodes a Di19 domain at its C-terminus, which has been implicated in recognition of MARylated histones (Longarini et al., 2023). Downstream of the Di19 domain is an annotated UIM, which has been shown previously to bind polyUb chains (Fisher et al., 2003; Giannini et al., 2008). Considering the proximity of these two domains, we hypothesized that they might cooperate to bind MARUbe modifications. We used AlphaFold3 to model the tandem Di19-UIM module at the C-terminus of RNF114 together with two Zn^2+^ ions, Ub, and NAD^+^ as a proxy for ADPr (Fig. 3A). The resulting model scored with high confidence across the entire complex, except the Ub C-terminus that showed indications of flexibility (Supplemental Fig. 3A-C). The top 5 models provided by AlphaFold3 were all highly similar. Interestingly, the models placed Ub in such an orientation that its C-terminus extended towards the adenine-proximal ribose of NAD^+^, where the 3’ hydroxyl group is the site of NAD^+^ and ADPr ubiquitylation by Deltex ligases. Upon closer inspection, there appeared to be a distinct pocket within the Di19 domain that bound the NAD^+^ moiety, with the nicotinamide group (which is replaced by a protein sidechain following ADP-ribosylation) exposed to solution and the only source of variation among the top 5 models. (Supplemental Fig. 3B). We identified two key coordination sites between the NAD^+^ and RNF114. Namely, W181 and G182 that are contained in a loop forming the back end of the NAD^+^-binding pocket, and R198 coordinating the diphosphate (Fig. 3B, Supplemental Fig. 3D). Y186 and E193 make additional hydrogen bonding contacts.

**Figure 3.**
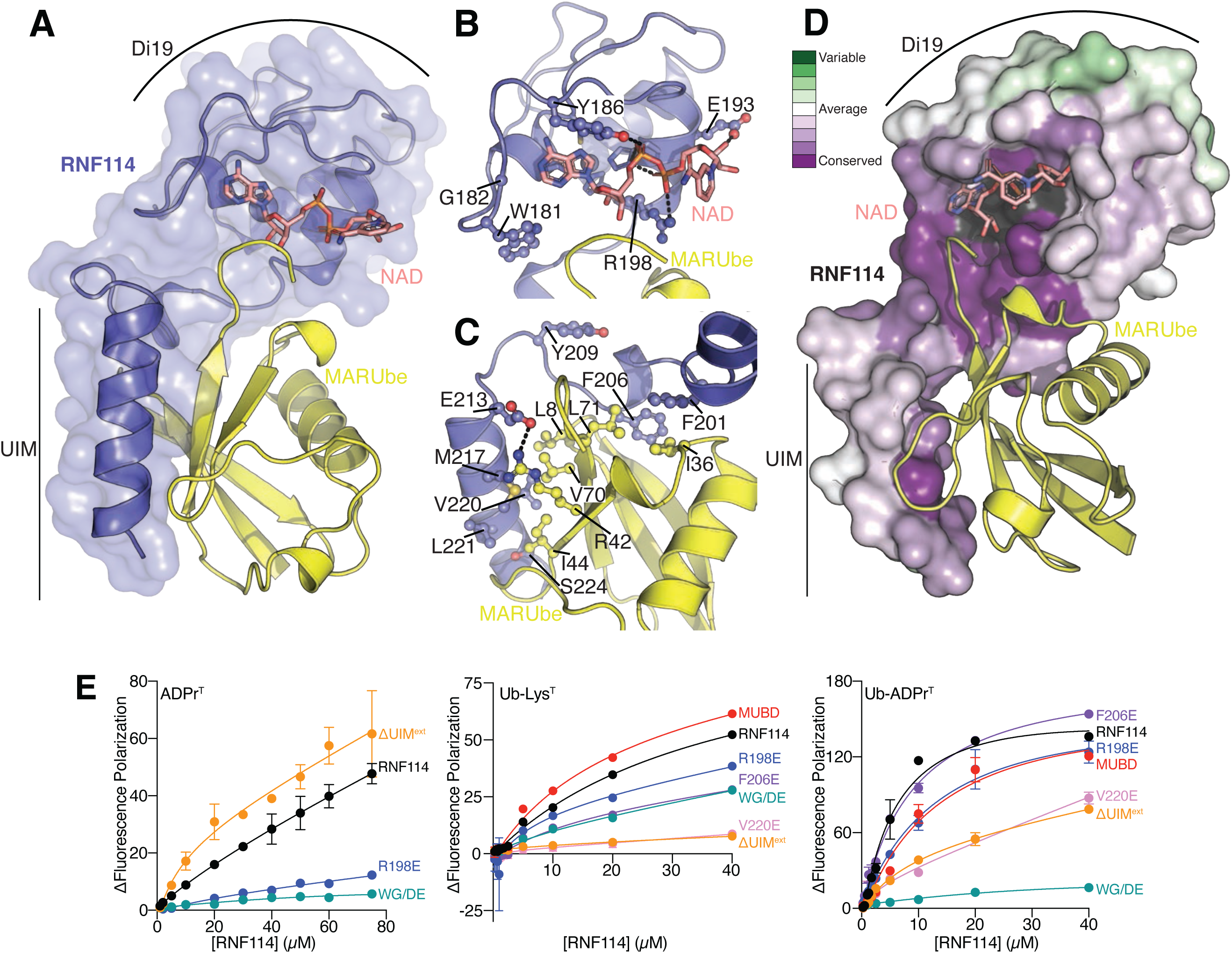
RNF114 binds ADPr and Ub. **A**) AlphaFold3 model of RNF114 Di19 and UIM domains (blue) in complex with two Zn^2+^ ions (grey spheres), as well as NAD (salmon) and Ub (yellow) to mimic a MARUbe modification. **B**) Detailed view of the modeled interaction between the RNF114 Di19 domain and NAD. **C**) Detailed view of the modeled interaction between the RNF114 UIM and Ub. **D**) Consurf analysis showing the conservation within the Di19 domain and UIM of RNF114 throughout evolution. As shown, dark purple represents the conserved areas, while green areas are more variable. The surface shown in dark grey represents the highly conserved Zn^2+^-binding structural residues in the Di19 domain. **E**) Model-guided mutations in RNF114 alter binding to ADPr^T^ (left), Ub-Lys^T^ (middle), and Ub-ADPr^T^ (right) substrates.

The C-terminal helix of RNF114 is a canonical UIM that coordinates Ub through the I44 patch (Fig. 3C). V220, centered in the UIM of RNF114 is a key conserved residue among UIMs. The UIM helix is also flanked on either end with highly conserved acidic residues and S224. The UIM is extended at its N-terminus to also coordinate the Ub I36 patch with F201/F206 of RNF114, located in the short linker region between its Di19 and UIM domains. When examining the evolutionary conservation of the RNF114 Di19-UIM module using ConSurf, we found strong conservation in the NAD^+^-binding site and on the Ub-interacting surface of the UIM helix (Fig. 3D). Our AlphaFold3 model therefore suggests that RNF114 is poised to recognize MARUbylation marks through coordination of its Di19 domain and C-terminal UIM. Thus, as these modeling studies suggest that the two domains work in concert, we propose to rename the tandem Di19-UIM module the <u>M</u>AR<u>U</u>be-<u>B</u>inding <u>D</u>omain (MUBD) and will refer to it as such going forward.

### MUBD mutants impair Ub-ADPr binding

To validate the AlphaFold3 model of the RNF114:NAD^+^:Ub complex, we rationalized point mutations of residues involved in binding ADPr, Ub, or Ub-ADPr. We tested these different RNF114 mutants for binding to our fluorescent substrates ADPr^T^ and Ub-ADPr^T^ (Fig. 3E, Supplemental Fig. 3E). As a control for a canonical Ub modification, we also compared to the Ub-Lys^T^ substrate generated previously (Geurink et al., 2012), in which Ub is isopeptide-linked to a TAMRA-labeled Lys-Gly dipeptide. Mutations within the ADPr-binding pocket, including W181D/G182E (herein referred to as WG/DE) or R198E, nearly eliminated the ability of RNF114 to bind ADPr^T^ (Fig. 3E, left; Supplemental Fig. 3E). A truncated version of RNF114, ΔUIM^ext^, which ends immediately after the Di19 domain at residue 200, retained ADPr^T^ binding comparable to WT RNF114. Though we could not reach saturation with our fluorescence polarization binding assay, we estimate the K_d_ of RNF114 for ADPr^T^ to be ∼40 µM.

Using Ub-Lys^T^ as a model for canonical Ub modifications, we could detect binding to RNF114 by fluorescence polarization but were again unable to calculate a K_d_ for the interaction due to the high concentrations of enzymes needed, though we estimate it to be ∼20 µM. We found that removal of the RING domain and associated disordered regions in RNF114 to create the MUBD construct (residues 137-228) facilitated slightly better Ub binding than full length RNF114 (Fig. 3E, middle; Supplemental Fig. 3E). The two ADPr-binding pocket mutants retained some of their ability to bind the Ub-Lys^T^ substrate. Surprisingly, the F206E version of RNF114, which disrupts coordination of the Ub I36 patch, retained some Ub binding. The ΔUIM^ext^ and V220E mutations significantly abrogated Ub binding, indicating the driving factor behind the noncovalent RNF114:Ub interaction lies at the center of the UIM.

Finally, we used Ub-ADPr^T^ to study the binding of RNF114 mutants to MARUbe (Fig. 3E, right; Supplemental Fig. 3E). Consistent with a multimode interaction, we detected much stronger binding to Ub-ADPr^T^ compared to ADPr^T^ and Ub-Lys^T^. For WT RNF114, binding was saturated by 40 µM, and we calculated a K_d_ of 12.8 µM. As observed with the ADPr^T^ or Ub-Lys^T^ substrates, the WG/DE, V220E, and ΔUIM^ext^ mutants of RNF114 were defective in their binding of Ub-ADPr^T^. Similar to the Ub-Lys^T^ substrate, the R198E mutant exhibited binding to Ub-ADPr^T^ at nearly the same level as WT RNF114, indicating that R198E can overcome the inability to bind ADPr whereas the WG/DE mutant could not. Another possibility is that R198 is not important for ADPr binding. Interestingly, we saw that RNF114 F206E retained binding of Ub-ADPr^T^ at similar levels as WT RNF114, while its binding to Ub-Lys^T^ was somewhat impaired. Together, these findings highlight the necessity of both the ADPr- and Ub-binding sites for a high-affinity interaction with MARUbe, but also that either binding site alone can maintain a weak affinity.

### RNF114 preferentially extends Ub-ADPr with polyUb

It is well accepted that many E3 ligases undergo rapid autoubiquitylation or catalyze the formation of free polyUb chains *in vitro*. We indeed observed free polyUb formation over the course of one hour with RNF114 and Ub, confirming our construct was active (Supplemental Fig. 4A). We then sought to use our fluorescent substrates in a UbiReal experiment (Franklin & Pruneda, 2019) to examine activity towards Ub-ADPr^T^ compared to Ub-Lys^T^, ADPr^T^, or Ub-Lys^T^ in the presence of ADPr (Fig. 4A). In these experiments, unlabeled Ub was added to facilitate ubiquitylation of the fluorescently labeled substrate. Under these conditions, the fluorescent Ub substrates can only behave as the “acceptor” during polyUb formation due to modifications at their C-termini, while unlabeled Ub serves as the “donor”. Formation of polyUb exclusively with unlabeled Ub is possible, but blind to the fluorescence polarization readout of the UbiReal experiment. In the presence of ligase reaction components, we observed no increase in fluorescence polarization of ADPr^T^ following addition of ATP, indicating that RNF114 cannot form the initial Ub-ADPr. Conversely, we observed a robust increase in fluorescence polarization with substrates containing Ub, indicating extension of polyUb products. Remarkably, RNF114 demonstrated a strong preference for extending the Ub-ADPr^T^ substrate compared to the Ub-Lys^T^ substrate, regardless of whether unlabeled ADPr was added to supplement the Ub-Lys^T^ reaction. This finding suggests that RNF114 recognizes the linkage between the Ub and ADPr moieties in MARUbe, and that simply occupying both binding sites with separate Ub and ADPr does not enhance ligase activity.

**Figure 4.**
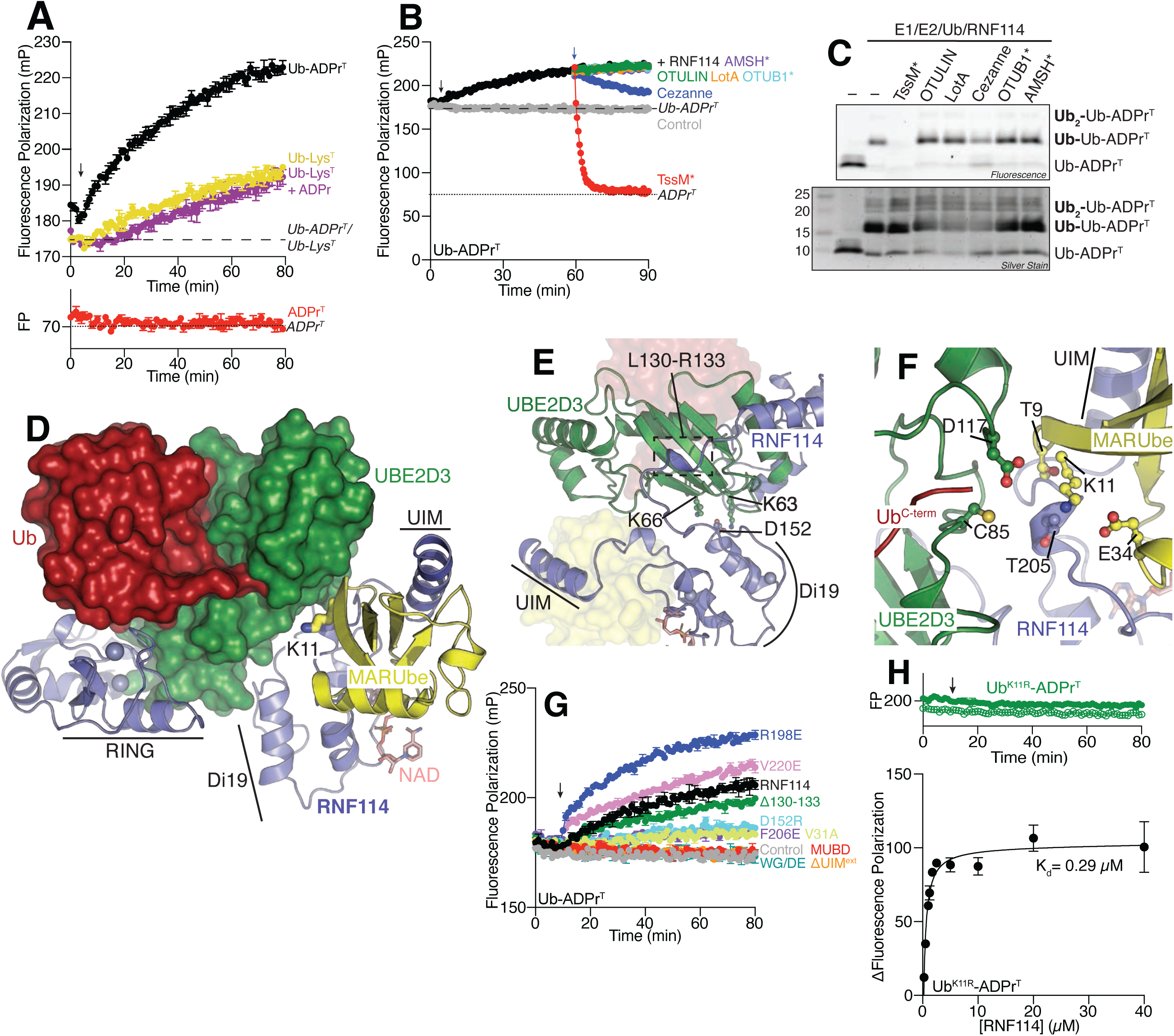
RNF114 extends K11 polyUb onto Ub-ADPr. **A**) UbiReal experiment showing the activity of RNF114 against the indicated substrates. The expected fluorescence polarizations of the substrates alone are indicated using dashed lines. Composition of the ligase experiment can be found in the Materials and Methods. **B**) The products of a ligase reaction were subjected to a UbiCRest experiment to identify the linkage type present on the Ub-ADPr^T^ substrate. Reactions were treated with the indicated deubiquitylase (added at the blue arrow), and their fluorescence polarization was monitored over time. After 30 minutes, the plate was removed from the instrument and samples were quenched in SDS sample buffer and **C**) fluorescent products were visualized by SDS-PAGE. **D**) AlphaFold3 model of the complex between RNF114 (blue), NAD (salmon), UBE2D3 (green), two copies of Ub (red/yellow), and five Zn^2+^ ions (grey spheres). Each protein and the important regions in RNF114 are labeled. **E**) RNF114 coordinates the E2 protein through backside binding (loop L130-R133) and charge interactions by D152. **F**) RNF114 positions the MARUbe so the K11 sidechain extends into the active site of UBE2D3∼Ub. To facilitate catalysis, a TEK box comprised of the MARUbe T9 and E34 are within proximity to deprotonate K11 of the MARUbe. RNF114 T205 is also in close proximity. **G**) A fluorescence polarization UbiReal experiment was used to evaluate the indicated regions of RNF114 and their potential roles in catalyzing the formation of Ub chains. **H**) A representative UbiReal experiment of a full ligase reaction (solid circles) for RNF114 with a K11R version of the Ub-ADPr^T^ (Ub^K11R^-ADPr^T^) substrate compared to the Ub^K11R^-ADPr^T^ substrate-only control (open circles). To examine binding, various concentrations of WT RNF114 were incubated with Ub^K11R^-ADPr^T^, and the data were collected as indicated in the Materials and Methods. The K_d_ was derived using GraphPad Prism 10 using a non-linear regression. Throughout all panels, the black arrow shows the point when ATP was added to the reaction.

### RNF114 specifically extends K11 polyUb on a Ub-ADPr substrate

Confirming that RNF114 exhibited preferential ligase activity toward Ub-ADPr^T^, we asked what type of Ub linkage RNF114 adds. To address this question, we used a UbiCRest experiment whereby the products of a ligase reaction (like those shown in Fig. 4A) were treated with a panel of deubiquitylases specific to distinct chain topologies (Hospenthal et al., 2015) (Fig. 4B). We selected deubiquitylases that are specific for Ub esters (TssM*; (Szczesna et al., 2024)), M1 polyUb (OTULIN;(Keusekotten et al., 2013)), K6 polyUb (LotA; (Warren et al., 2023)), K11 polyUb (Cezanne; (Mevissen et al., 2016)), K48 polyUb (OTUB1*; (Michel et al., 2015)), or K63 polyUb (AMSH*; (Michel et al., 2015)). As we had previously observed K11 extension of MARUbe on PARP7 and PARP10 from cells (Bejan et al., 2025), and observed reduced polyUb extension on PARP7 MARUbe following RNF114 knockdown (Figure 1D), we hypothesized that RNF114 was extending K11 polyUb onto Ub-ADPr. Initial testing of Cezanne against the RNF114 ligase products demonstrated that high concentrations of this enzyme were able to cleave the Ub-ADPr ester linkage, as evident by the fluorescence polarization returning to values <100, consistent with the observed fluorescence polarization of ADPr^T^ (Supplemental Fig. 4B). We therefore optimized a concentration of 20 nM Cezanne, which did not cleave the Ub-ADPr ester linkage after 30 minutes. Importantly, all of the other polyUb deubiquitylases included in our UbiCRest panel are exclusive to their polyUb substrates and cannot cleave other types of Ub modification (Hospenthal et al., 2015; Warren et al., 2023). The UbiCRest experiment demonstrated that the only deubiquitylase other than Cezanne that cleaved RNF114 ubiquitylation products was the Ub esterase TssM*. Notably, Cezanne treatment returned the fluorescence polarization to ∼180, in good agreement with values for Ub-ADPr^T^ alone (Fig. 4B). Treatment with TssM* returned the fluorescence polarization to ∼70, in good agreement with the observed polarization of ADPr^T^ alone, indicating that TssM* had likely cleaved the entire polyUb chain from ADPr. Indeed, when we examined products of the UbiCRest experiment by SDS-PAGE and in-gel fluorescence, we observed strong signal for Ub-Ub-ADPr^T^ produced by RNF114 (Fig. 4C). In the presence of TssM*, the fluorescence signals corresponding to Ub-Ub-ADPr^T^ and Ub-ADPr^T^ were eliminated, consistent with release of ADPr^T^. Upon silver staining, we observed an intense protein band correlating to diUb, confirming our suspicions that TssM* had cleaved the entire polyUb from ADPr^T^. Cezanne treatment decreased the intensity of the Ub-Ub-ADPr^T^ band by ∼50%, and we observed the reappearance of a fluorescent band at the molecular weight of Ub-ADPr^T^. The other tested deubiquitylases showed no decrease in fluorescence of the Ub-Ub-ADPr^T^ species. We then completed the same UbiCRest experiment for ligase products using Ub-Lys^T^ to address whether specificity was retained (Supplemental Fig. 4C). For products generated with Ub-Lys^T^, we again observed robust cleavage with Cezanne, indicating the specific formation of K11 polyUb. As expected in this experiment, TssM* did not cleave the products, as there was no ester-linked Ub present in the reaction. Taken together, these findings support the hypothesis that RNF114 specifically extends K11 polyUb on its preferred target of Ub-ADPr.

### Molecular basis for RNF114 extension of K11 polyUb on Ub-ADPr

To examine the molecular basis for K11 extension of MARUbe by RNF114, we used AlphaFold3 to model the full transferase complex containing full length RNF114, UBE2D3, NAD^+^, two copies of Ub, and five Zn^2+^ ions (Fig. 4D). Remarkably, the model had high confidence throughout most of the complex (Supplemental Fig. 4D, E). Outside of a disordered region N-terminal to the RNF114 RING domain, only one region of RNF114, between residues 128-134 that cross over the backside of the E2 protein, was scored as low confidence. The top 5 scoring models from AlphaFold3 aligned nearly perfectly, with slight variation at the N-terminus of RNF114 and in the positioning of the NAD^+^ nicotinamide group (Supplemental Fig. 4F). We observed a canonical E2:RING interface whereby helix α1, and loops 4 and 7 of UBE2D3 mediate the interaction (Supplemental Fig. 4G). The RING:E2 interface is extended by an additional RNF114 zinc finger that binds UBE2D3 helix 1 analogously to a previously determined crystal structure of RNF125 (Middleton et al., 2023). We also observed UBE2D3 backside coordination by RNF114 residues 128-134 and a potential charge interaction by D152 (Fig. 4E). One copy of Ub was found to mimic a thioester intermediate with UBE2D3, wherein the Ub C-terminus extended toward the UBE2D3 C85 active site, and the two proteins formed a ‘closed’ conformation of E2∼Ub observed in RING-dependent priming of Ub transfer (Dou et al., 2012; Plechanovova et al., 2012; Pruneda et al., 2012). The second copy of Ub, together with NAD^+^, bound into the MUBD module at the RNF114 C-terminus in a manner identical to our previous modeling. Consistent with our biochemical analysis of RNF114 specificity, further examination of this complex revealed the K11 sidechain of the MARUbe was around 6 Å away from and pointed directly towards the active site of UBE2D3, supported by residues from both the E2 and RNF114 (Fig. 4F). Thus, the full transferase complex model suggests K11-specific ubiquitylation of MARUbe. We designed a series of point mutations to validate the model, which were tested in our UbiReal experiments (Fig. 4G, Supplemental Fig. 4H, I). As expected, none of the RNF114 mutants tested could ubiquitylate the ADPr^T^ substrate, and many mutants showed no activity towards Ub-Lys^T^ either in the absence or presence of unlabeled ADPr (Supplemental Fig. 4I). We observed ubiquitylation of the Ub-Lys^T^ substrate with the R198E or Δ130-133 versions of RNF114, suggesting that 1) Ub binding to the UIM can overcome charge differences in the ADPr-binding pocket; and 2) the backside binding of UBE2D3 mediated by residues 130-133 of RNF114 does not influence ubiquitylation. Other E2-coordinating mutants, V31A or D152R were completely defective in their ubiquitylation of Ub-Lys^T^. All E2-binding mutants of RNF114 (V31A, Δ130-133, and D152R) retained some ubiquitylation of the Ub-ADPr^T^ substrate, where the Δ130-133 mutant exhibited ligase activity comparable to WT RNF114, and the V31A and D152R mutants reduced activity (Fig. 4G). The F206E mutant of RNF114 displayed ubiquitylation activity similar to V31A and D152R. Surprisingly, we observed robust ubiquitylation with the V220E and R198E mutants of RNF114, suggesting that despite compromised binding of either the ADPr^T^ or Ub-Lys^T^ substrates, RNF114 ligase activity does not require tight binding of the MARUbe to facilitate its ubiquitylation. There were three mutants that did not possess any ligase activity towards the Ub-ADPr^T^ substrate: WG/DE, MUBD, and ΔUIM^ext^. The lack of activity was expected for the WG/DE and MUBD mutants due to their inability to bind Ub-ADPr^T^ and the lack of the RING domain, respectively. However, we were intrigued to see that the ΔUIM^ext^ version of RNF114 failed to ubiquitylate the Ub-ADPr^T^ substrate despite its only moderately impaired binding of the substrate (Fig. 3E, right). It is possible that, while these mutants can still bind Ub-ADPr, an interaction within the UIM is required to orient Ub into the E2 active site. Taken together, these data not only validate the structural model but also show that RNF114 ligase activity can overcome defective binding due to mutation in either the Ub or ADPr binding sites.

Because the structural modeling, coupled with our earlier UbiCRest data, suggested tight specificity for K11 ubiquitylation, we sought to test whether RNF114 could produce any other polyUb product. To address this, we used a K11R mutant of Ub and the methodology described in Fig. 2 to form Ub^K11R^-ADPr^T^ (Supplemental Fig. 4J). We validated the binding of this substrate to WT RNF114 and were surprised to see much tighter binding compared to the Ub-ADPr^T^ substrate (K_d_= 290 nM) (Fig. 4H). In a UbiReal experiment, RNF114 failed to extend any Ub onto the Ub^K11R^-ADPr^T^ substrate, even after one hour (Fig. 4H). This result again confirms that RNF114 specifically extends K11 linkages on the Ub-ADPr substrate. Furthermore, none of the structure-guided RNF114 mutants could extend Ub onto the Ub^K11R^-ADPr^T^ substrate (Supplemental Fig. 4K), confirming that none of the retained activity observed with some mutants was due to formation of different polyUb linkages. Taken together, these data further indicate that RNF114 synthesizes exclusively K11 linkages on a Ub-ADPr substrate.

### RNF114 belongs to a family of MARUbe-Targeted Ligases (MUTLs)

The initial identification of RNF114 as a Di19 domain-containing protein also identified Di19 domains in the E3 ligases RNF125, RNF138, and RNF166 (Fig. 5A) (Giannini et al., 2008). A sequence alignment of these Di19-containing E3 ligases showed a conservation of the Zn^2+^- and E2-coordinating residues, as well as MUBD architecture (Supplemental Fig. 5A). One key difference we noticed was the length of the linker connecting the RING-ZnF ligase module to the MUBD, where RNF114 and RNF166 had the same linker length, while the linkers in RNF125 and RNF138 were shorter or longer, respectively. Interestingly, we observed a conserved ADPr-binding site in RNF114, RNF138, and RNF166 consisting of a WGD motif, which was missing in RNF125. Specifically, the residues that create part of the backside of the ADPr-binding pocket (W181/G182) were not conserved in RNF125. Instead, RNF125 possesses D178/E179, which based on our mutagenesis work with RNF114, diminish ADPr binding. Key residues in the UIM region remain conserved, including the charged residues on the N-terminal side of the helix and the polar serine near the C-terminus.

**Figure 5.**
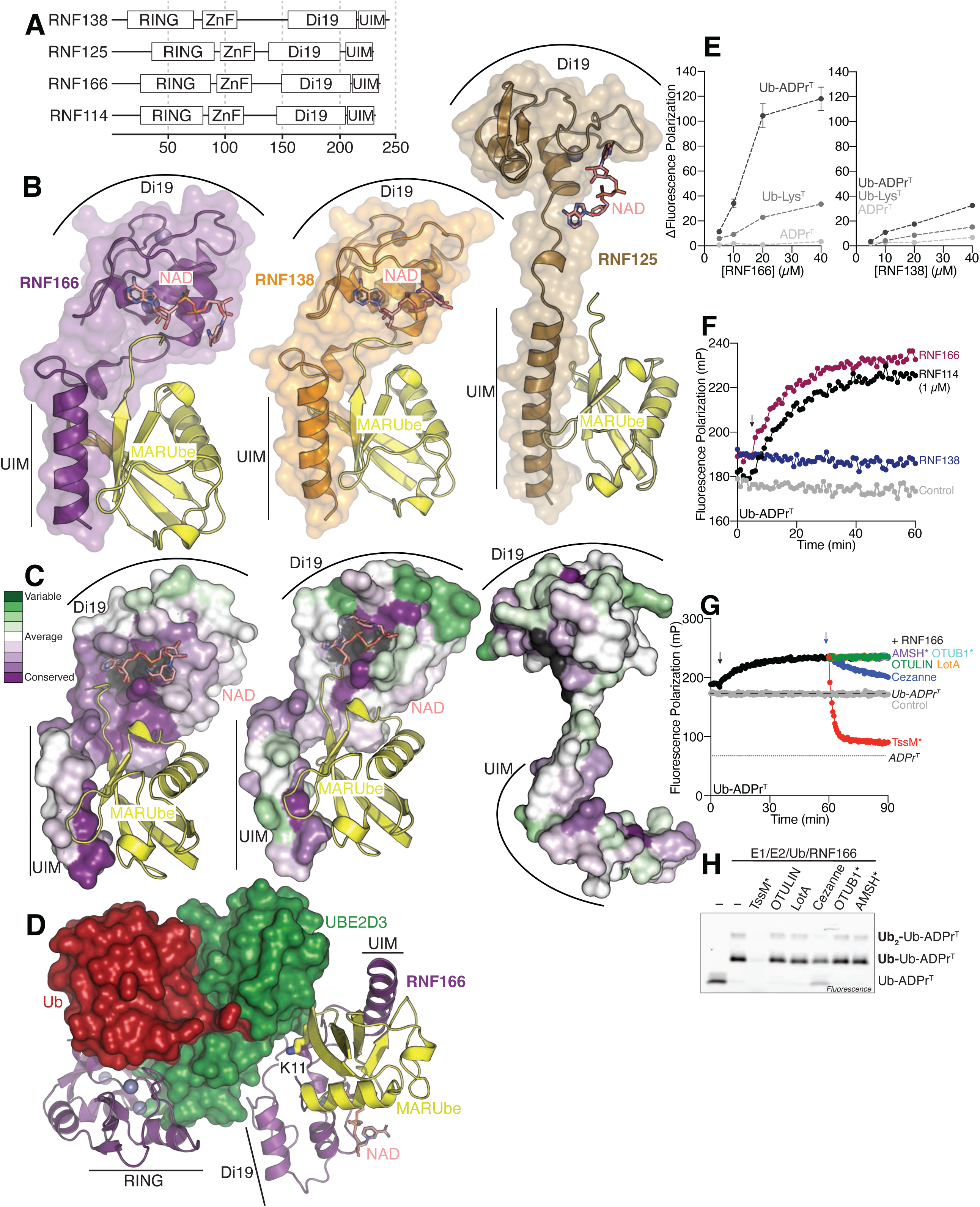
RNF114 belongs to a family of <u>M</u>AR<u>U</u>be-<u>T</u>argeted <u>L</u>igases (MUTLs). **A**) Schematic representation of the domain organization for RNF114, RNF125, RNF138, and RNF166, other Di19 domain- and UIM-containing E3 ligases. **B**) AlphaFold3 models depicting the MUBD of the indicated E3 ligases in complex with two Zn^2+^ ions, as well as NAD and Ub to mimic MARUbe. **C**) Consurf analysis of the indicated E3 ligases ordered as above showing evolutionary conservation (purple) or variability (green). **D**) AlphaFold3 model of RNF166 (violet), UBE2D3 (green), NAD (salmon), two copies of Ub (red/yellow), and five Zn^2+^ ions (grey spheres). Shown is the positioning of the MARUbe K11 sidechain extending towards the UBE2D3 active site in an orientation similar to the model for RNF114. **E**) RNF166 and RNF138 preferentially bind Ub-ADPr^T^ over Ub-Lys^T^ and ADPr^T^ substrates. **F**) Representative UbiReal experiment for the indicated E3 ligases with the Ub-ADPr^T^ substrate. The control represents the baseline fluorescence polarization of the substrate without any other proteins. ATP was added to all reactions at the timepoint indicated by the black arrow. In this experiment, RNF114 is 1 µM, and the other ligases are 5 µM. **G**) The products of an RNF166 ligase reaction (black dataset) were subjected to a UbiCRest experiment with the indicated deubiquitylases added at 60 minutes, indicated by the blue arrow. **H**) Products of the UbiCRest were evaluated using in-gel fluorescence following SDS-PAGE. Throughout the panels, the black arrow marks the point when ATP was added to the reactions.

Based on this sequence alignment, we wondered if these other ligases would show activity for a MARUbe substrate. We used AlphaFold3 to model RNF125, RNF138, or RNF166 with NAD^+^ and Ub to evaluate possible binding to the MUBD (Fig. 5B). For RNF138 and RNF166, which had higher similarity with RNF114, AlphaFold3 modeled the complexes with high confidence (Supplemental Fig. 5B). Consistent with our model of RNF114 (Fig. 3, Supplemental Fig. 3), the nicotinamide group was observed in several orientations extending into solution (Supplemental Fig. 5C). Consistent with the lack of conservation in the ADPr-binding pocket, AlphaFold was unable to produce a high confidence model for RNF125 in complex with NAD^+^ and Ub. When examined with ConSurf, RNF166, RNF138, and RNF125 each had a different level of evolutionary conservation (Fig. 5C). Notably, RNF166 possessed a nearly continuous conserved patch stretching from the ADPr-binding site to the Ub-binding face of the UIM. RNF138 demonstrated regions of conservation in the ADPr-binding site and toward the C-terminus of the UIM with regions of variation dispersed throughout the MUBD. RNF125, however, demonstrated little conservation including throughout the ADPr-binding site, suggesting that RNF125 might not bind NAD^+^ (Supplemental Fig. 5B, C). We then selected RNF166 to model a full transferase complex in AlphaFold3 (Fig. 5D) and again observed the MARUbe K11 sidechain extending towards the active site of UBE2D3∼Ub. We also noticed that the model exhibited high confidence throughout the majority of the complex and the top 5 scoring models were very similar (Supplemental Fig. 5D, E).

We aimed to validate the AlphaFold3 models with Ub-ADPr binding experiments for each of the three ligases; however, we could not produce soluble RNF125 for these experiments. Similar to RNF114, and as expected from the AlphaFold3 models, both RNF166 and RNF138 showed selective binding for Ub-ADPr^T^ over ADPr^T^ and Ub-Lys^T^, as demonstrated by much higher changes in fluorescence polarization (Fig. 5E). RNF166 appeared to have similar binding affinity for Ub-ADPr^T^ as RNF114, while RNF138 did not bind Ub-ADPr^T^ as well. These binding assays corroborate the AlphaFold3 models and sequence alignment, showing that RNF114, RNF166, and RNF138 all have functional MUBD domains capable of reading MARUbe modifications.

We next evaluated the ligase activity of RNF138 and RNF166 in a UbiReal experiment with our fluorescent substrates. Surprisingly, only RNF166 was able to extend polyUb onto Ub-ADPr^T^ but required 5-times higher concentration than RNF114 (Fig. 5F). RNF166 was also able to ubiquitylate Ub-Lys^T^ at these higher concentrations, though we observed a strong preference for the Ub-ADPr^T^ substrate (Supplemental Fig. 5F). We performed a UbiCRest analysis to determine which chain topology RNF166 was building onto Ub-ADPr^T^ (Fig. 5G). Upon treating the ligase reaction with our panel of specific deubiquitylases, we observed that only TssM* and Cezanne were able to cleave the ubiquitylated Ub-ADPr^T^, similar to RNF114. TssM* cleaved the products down close to the expected polarization of ADPr^T^ (∼85), as shown by the signal decrease below the Ub-ADPr^T^ control. Cezanne, the K11-specific deubiquitylase, only partially cleaved the extended Ub-ADPr^T^, returning the polarization to ∼200. These samples were then evaluated on SDS-PAGE with in-gel fluorescence and we observed that the main product was Ub-Ub-ADPr^T^ and a minor amount of Ub_2_-Ub-ADPr^T^ (Fig. 5H). We also conducted a UbiCRest experiment on a RNF166 ligase reaction utilizing Ub-Lys^T^ as a substrate and observed the same trends as RNF114 (Supplemental Fig. 5G, H). Here, RNF166 synthesized longer linkages on the Ub-Lys^T^ substrate that were only cleaved by the K11-specific Cezanne. Taken together, we show that MUBD-containing E3 ligases can be considered a unique family of <u>M</u>AR<u>U</u>be-<u>T</u>argeted Ub <u>L</u>igases, or MUTLs, that can read MARUbe modifications and write new extensions of polyUb which, at least for RNF114 and RNF166, are specifically linked via K11.

## Discussion

Here, we show that a family of E3 Ub ligases containing a tandem Di19-UIM module can specifically recognize ubiquitylated mono-ADPr (MARUbe). We extensively evaluate the structure and function of one of these family members, RNF114, and decipher the mechanism by which RNF114 builds a K11-linked polyUb onto MARUbe. We show that RNF114 has a strong preference for the MARUbe substrate compared to ADPr or Ub alone (Fig. 3E, Fig. 4A). We highlight how RNF114 positions MARUbe so that K11 reaches towards the active site of an incoming E2∼Ub conjugate (Fig. 4D, F), and we extend our observations of RNF114 to a family of Ub ligases containing a tandem Di19-UIM. We found that at least 2 other members (RNF138, RNF166) preferentially bind MARUbe compared to ADPr or Ub alone, and one, RNF166, catalyzes the K11-linked ubiquitylation of MARUbe. We therefore suggest renaming the tandem Di19-UIM the <u>M</u>ar<u>U</u>be-<u>b</u>inding <u>d</u>omain (MUBD) and the family of ligases <u>M</u>AR<u>U</u>be-targeted ligases (MUTLs).

Although all members of the MUTL family possess a MUBD, we only observed ubiquitylation of Ub-ADPr with RNF114 and RNF166, suggesting differences in catalytic activity between each of the family members. One obvious source of variation is the linker length between the MARUbe reader and writer modules. RNF114 and RNF166 have the same linker length, while RNF138 is longer and RNF125 is shorter. The linker length could directly relate to the conformation of the writer and reader modules after MARUbe binding, whereby only the RNF114 and RNF166 complexes have the correct structural organization to facilitate MARUbe ubiquitylation. Interestingly, RNF125 is the only related ligase that does not retain a conserved MUBD (Fig. 5C), which could suggest that either the RNF125 MUBD is degenerate or that it is rapidly evolving toward a substrate other than MARUbe.

We and others have shown that various PARP proteins (PARP7, PARP10, Tankyrase) are MARUbylated in cells, and that the MARUbe modification is extended with K11 polyUb (Bejan et al., 2025; Perrard et al., 2025). A current model suggests that MARUbylation could stabilize a substrate by blocking PAR extension and preventing downstream PARdU, but it remains unclear if this outcome is consistent across all MARUbylated substrates (Perrard et al., 2025). PARPs are known to MARylate the sidechain of various residues including Glu/Asp, Ser/Thr, and Cys, and the lability of the bond changes with each. For PARP10 and now PARP7, we have shown that MARUbylation occurs specifically on Glu/Asp residues (Fig. 1A) (Bejan et al., 2025). Meanwhile, Tankyrase was suggested to be MARUbylated on serine (Perrard et al., 2025). Together, these findings suggest the lability and transient nature of MARUbylation, indicating the species might serve as an intermediate in some undescribed signaling pathway.

The abundant crosstalk between PARPs and ubiquitylation at sites of DNA damage provides an obvious axis for therapeutic intervention. PARP inhibitors such as olaparib, niraparib, rucaparib, and talazoparib are currently being used to treat various cancers (Del Campo et al., 2019; Matulonis et al., 2016; Moore et al., 2018; Swisher et al., 2017), and the natural product nimbolide has been shown to target RNF114, preventing its ubiquitylation and degradation of PARP1 (Li et al., 2023). The multi-layered regulation of a substrate through MARUbylation provides an additional opportunity for therapeutic development and the possibility of co-treatment with current PARP inhibitors. As MUTL activity occurs downstream of PARP activity, targeting the MUTL family of E3 ligases through their ability to bind and read marks of MARUbylation will provide an added layer of specificity unreachable solely with PARP inhibitors.

Overall, the work presented here structurally and functionally demonstrate specific K11 polyUb extension of MARUbe modifications by a family of related E3 ligases. Additional layers of regulation for this complex PTM, including at the level of MUTL activity, will be an interesting area of future work. Within RNF114, for example, there is strong evidence for phosphorylation at Tyr116, Tyr186, and Tyr209, which lie at interfaces with the E2, ADPr, and Ub, respectively (Fig. 3B, C, Supplemental Fig. 4G) (Hornbeck et al., 2015). Furthermore, the functional rationale for K11 polyUb specificity remains a fascinating mystery. While K11 linkages have previously being implicated in cell cycle regulation by the APC/C, they are in the context of heavily branched K11/K48 polyUb chains that accelerate proteasomal degradation (Wu et al., 2010; Yau et al., 2017). Alongside PARdU, MARUbylation represents a complicated component of the PARP/Ub axis, and future work will expand how these heavily regulated PTMs synergize in cellular signaling.

## Materials and Methods

### Cell culture

Cells were cultured at 37 °C and 5% CO_2_. HEK 293T (ATCC, CRL-3216), HEK 293 control and PARP1 KO cells lines (gift from Prof. Michael Garabedian at NYU Langone) were cultured in DMEM (Gibco, 11965118) supplemented with 10% FBS (Sigma-Aldrich, F0926), 1X GlutaMAX (Gibco, 35050061), and 1 mM sodium pyruvate (Gibco, 11360070).

### siRNA knockdown and GFP-PARP7 overexpression

HEK 293T cells were reverse transfected with 50 nM total siRNA (Horizon Discovery) targeting RNF114 (cat no. L-007024-00-0005), RNF166 (cat no. L-007119-00-0005), or a non-targeting control (cat no. D-001810-10-05), using DharmaFECT 1 Transfection Reagent (Horizon Discovery, cat no. T-2001-02). After 48 hr, cells were then co-transfected with GFP-PARP7 C5 and HA-Ub plasmids using jetOPTIMUS® DNA transfection Reagent (Polyplus-transfection, cat no. 101000025) for 24 hr (media was exchanged 4 hr post-transfection). RBN2397 was used at 300 nM for 18 hours following the media swap. Cells were then lysed and immunoprecipitated as described below.

### GFP immunoprecipitation for on-bead enzyme/chemical treatment assay (MARUbylation assay)

GFP-Trap® Magnetic beads (Proteintech, gtma) were washed twice with cell lysis buffer (CLB; 50 mM HEPES pH 7.4, 150 mM NaCl, 1 mM MgCl_2_, 1% Triton X-100) and added to lysates at ∼300-500 µg total protein/10 µl bead slurry. The beads were rotated for 2 hr at 4 °C, then washed once with CLB, twice with 7 M Urea/1% SDS in PBS, once with 1% SDS in PBS, and three times with HEPES buffer (Hb: 50 mM HEPES pH 7.5, 100 mM NaCl, 4 mM MgCl_2_, 0.2 mM TCEP added fresh). The beads were then treated as described below.

In Figure 1A, the beads were treated with TssM (2 µM), TssM* (2 µM), NUDT16 (10 µM, containing 15 mM MgCl_2_), or hydroxylamine (1 M, in HEPES buffer pH 7.5) for 1 hr at 37 °C. The supernatant was then separated from the beads and both fractions were quenched with 4X sample buffer (1X: 10% glycerol, 50 mM Tris-HCl (pH 6.8), 2% SDS, 1% β-mercaptoethanol, 0.02% bromophenol blue). The bead fraction was boiled at 95 °C for 5 min to elute GFP-PARP7 for detection by western blotting.

In Figure 1D, the beads were treated with NUDT16 (10 µM, containing 15 mM MgCl_2_) for 1 hr at 37 °C. The supernatant was then separated from the beads and both fractions were quenched with sample buffer. The bead fraction was boiled at 95 °C for 5 min to elute GFP-PARP7 for detection by western blotting.

### Plasmids and cloning

DNA sequences for human RNF114, RNF138, and RNF166 were synthesized from Twist Biosciences and subsequently cloned into the pOPIN-B and pOPIN-S3C vectors (Berrow et al., 2007). Site-directed mutagenesis was used to insert substitutions into RNF114 and products were confirmed with DNA sequencing.

### Protein expression

The following proteins were purified as previously reported: Human UBE1 (Gladkova et al., 2018), UBE2D3/DTX2 (Bejan et al., 2025), AF1521, NUDT16, TssM* (Szczesna et al., 2024), OTULIN (Keusekotten et al., 2013), LotA-N/Ub (Warren et al., 2023), Cezanne (Mevissen et al., 2016), OTUB1*/AMSH* (Michel et al., 2015).

His-tagged RNF114 and RNF166 were expressed as N-terminal His-3C fusion proteins from the pOPIN-B vector and RNF138 was expressed as an N-terminal His-SUMO-3C fusion from pOPIN-S3C using fresh transformations of *E. coli* Rosetta cells. A single colony was used to inoculate a starter culture of selective Luria Broth (LB) (30 µg/mL kanamycin, 35 µg/mL chloramphenicol). The starter was used to inoculate larger LB cultures, which were grown at 37 °C until the OD_600_ reached 0.4-0.6. 50 µM ZnCl_2_ was added just before induction with 0.2 mM isopropyl β-D-1-thiogalactopyranoside (IPTG) for overnight expression at 18 °C. The cells were harvested by centrifugation at 4500g, and in the case of RNF114 and RNF166, pellets were resuspended in 25 mM Tris pH 7.4, 200 mM NaCl, 2 mM β-mercaptoethanol. Sigmafast protease inhibitor cocktail, PMSF, lysozyme and DNase were added to the cell solutions prior to lysis by sonication. The lysate was clarified by centrifugation at 45000g, at 4 °C for 40 minutes. The supernatant containing the His-tagged protein was applied to Cobalt resin, and the resin was washed extensively with 25 mM Tris pH 7.4, 200 mM NaCl, 2 mM β-mercaptoethanol. Bound proteins were eluted using the same buffer containing 250 mM imidazole. Eluted proteins were further purified using gel filtration chromatography on a HiLoad 16/600 Superdex75 in 25 mM Tris pH 7.4, 150 mM NaCl, 2 mM DTT. RNF138 was purified using the same protocol with the exception that buffers were at pH 8.0. Fractions containing the protein of interest were pooled, concentrated, flash frozen, and stored at - 70°C until use.

### Synthesis of clickable ADPr (Compound 8)

Methods and validation corresponding to the synthesis of our clickable ADPr analog can be found in the supporting information.

### Preparation of Ub-ADPr-TAMRA (Ub-ADPrT)

Ub-ADPr was generated similar to previously described using DTX2 and our clickable ADPr analog (Bejan et al., 2025; Kelly et al., 2024; Zhu et al., 2024; Zhu et al., 2022). Briefly, a preparative amount of E2∼Ub conjugate was formed using either WT Ub or Ub^K11R^ and purified using a Superdex75 10/300 Increase column pre-equilibrated in 25 mM Hepes pH 7.4, 150 mM NaCl. This step is necessary to remove any remaining ATP or its hydrolyzed forms that have previously been shown to be alternative substrates of DTX2 (Zhu et al., 2022). Any remaining compound in the ubiquitylation reaction that contains an ADP-ribose would be ubiquitylated by DTX2, resulting in heterogenous products instead of homogenous Ub-ADPr. The purified conjugate was then mixed with 5 µM DTX2 and 0.2 mM compound **8** in 25 mM Hepes pH 7.4, 150 mM NaCl to enzymatically form Ub-ADPr. After 1 hour at 37 °C, the reaction was diluted 1:10 in 25 mM NaOAc pH 4.5. Ub-ADPr was purified from other reaction components using cation exchange chromatography on a Resource S column over a gradient from 0-500 mM NaCl. Fractions corresponding to Ub-ADPr were pooled and concentrated to <150 µL and evaluated using intact mass spectrometry.

The fluorescent species, Ub-ADPr^T^, was made using click chemistry between the ADPr (alkyne) of Ub-ADPr and the TAMRA molecule (azide). A 3X reaction mixture was made up using 1.5 mM Tris(3-Hydroxypropyltriazolylmethyl)amine (THPTA), 0.75 mM CuSO_4_·5H_2_O, 0.3 mM TAMRA-azide, and 7.5 mM sodium ascorbate in PBS pH 7.4. This mixture was then diluted to 1X into the Ub-ADPr solution and mixed for 1.5 hours at room temperature. Ub-ADPr^T^ was separated from unconjugated TAMRA-azide using a HiLoad 16/600 Superdex75 size exclusion column pre-equilibrated in 25 mM NaOAc pH 4.5, 150 mM NaCl. Fractions corresponding to Ub-ADPr^T^ were pooled and concentrated, and the concentration was measured using the TAMRA fluorophore. The stock of Ub-ADPr^T^ was diluted back to more neutral pH using 50 mM Hepes pH 7.4, 100 mM NaCl prior to long term storage at -70 °C.

### Gel-based ubiquitylation experiment

Ubiquitylation experiments were conducted at 37°C with 0.2 µM E1, 5 µM UBE2D3, 50 µM Ub, 2 µM RNF114, 1 mM ATP, 5 mM MgCl_2_, and 1 mM DTT in 50 mM HEPES, pH 7.4, and 100 mM NaCl. The reaction was quenched with sample buffer at the indicated timepoints and resolved on 4/16.5% Tris-Tricine gel. The gel was stained with Coomassie blue to visualize ubiquitylation products.

### Fluorescence polarization experiments

Fluorescence polarization experiments were conducted using a BMG Labtech CLARIOstar microplate reader. Reaction volumes were 10 µL and data were collected in a black, low protein-binding 384-well plate at 22 °C. For all experiments, 25 mM Hepes pH 7.4, 150 mM NaCl, 0.1 mg/mL BSA, 1 mM TCEP (FP buffer) was used as the blank. The 540-20 nm and 590-20 nm optic filters were used to monitor the TAMRA fluorophore.

For binding experiments, a range of concentrations were chosen for each enzyme. Fresh dilutions were made in duplicate using FP buffer. Samples were mixed 1:1 using a 2X stock of the indicated fluorescent substrate (Ub-ADPr^T^, ADPr^T^, Ub-Lys^T^) and protein. Ten measurements were recorded per sample, and an average was taken. To determine K_d_ values of RNF114 with Ub-ADPr^T^ or Ub^K11R^-ADPr^T^ and AF1521 with ADPr^T^, the average baseline signal of the fluorescent substrate in the absence of binding partner was subtracted from each measurement and final values were plotted as change in fluorescence polarization (ΔFP) against protein concentration. A non-linear regression on the mean ± SEM of the duplicate datasets was used in GraphPad Prism 10 to derive K_d_ values with the ADPr^T^, Ub-ADPr^T^, Ub^K11R^-ADPr^T^ substrates.

A UbiReal experiment was used to monitor the ligase activity of proteins (Franklin & Pruneda, 2023). A reaction mixture containing 0.2 µM E1, 1 µM UBE2D3, 10 µM unlabeled Ub, 0.5 µM RNF114, 5 mM MgCl_2_, with 50 nM of the indicated fluorescent substrate (Ub-ADPr^T^, ADPr^T^, Ub-Lys^T^) was made up in FP buffer. For ligase reactions with RNF138 and RNF166, 5 µM ligase was used. A baseline FP was collected for 10 minutes and then 1 mM ATP was added to the wells to initiate the ligase assay. The plate was put back into the plate reader and monitored for 70 more minutes. For reactions containing unlabeled ADPr, 0.2 mM was used.

For the UbiCRest experiment to diagnose the chain type formed by RNF114, the products of a UbiReal experiments were treated with TssM*, OTULIN, LotA, OTUB1*, or AMSH* at 0.5 µM concentration or with Cezanne at 0.02 µM concentration. The reaction was monitored by fluorescence polarization for the indicated amount of time. At the end of the reaction, samples were removed from the plate, quenched with sample buffer and resolved on a 4-20% Mini-PROTEAN TGX gel (Bio-Rad Laboratories). Reaction products were visualized by in-gel fluorescence to track TAMRA-conjugated species.

### Mass spectrometry

Intact mass was determined using a ThermoScientific LTQ Velos Pro linear ion with a C18 trap. A Poroshell C18 column (1.0 Å x 75 mm x 5 µm) was used for reverse-phase high performance liquid chromatography (RP-HPLC) prior to sample ionization. 0.5 µg of purified protein was resolved using a water/acetonitrile gradient from 2-50% acetonitrile in 0.1% formic acid. Data were acquired using Xcalibur (ThermoScientific) and processed using Freestyle (ThermoScientific) for spectrum deconvolution.

### Western blotting

Cells were lysed in cell lysis buffer (CLB; 50 mM HEPES pH 7.4, 150 mM NaCl, 1 mM MgCl2, 1% Triton X-100) supplemented with fresh 1 mM TCEP, 1X cOmplete EDTA-free Protease Inhibitor Cocktail (Sigma-Aldrich, 11873580001), 30 μM Phthal 01 (pan-PARP inhibitor) (Rodriguez et al., 2021), and 50 µM PR-619 (Medchemexpress, HY-13814). Lysates were centrifuged at 14,000 rpm for 10 min at 4 °C, quantified with a Bradford assay (Bio-Rad Laboratories, 5000006EDU), and resolved in 4–20% precast gels (Bio-Rad Laboratories, 4561096). Proteins were transferred to a nitrocellulose membrane using Trans-Blot Turbo Transfer System (Bio-Rad Laboratories), blocked in 5% milk (Carnation) in 1X PBS, 0.1% Tween-20 (PBST), and probed overnight at 4 °C with primary antibodies. After three washes with PBST, the blots were incubated for 1 hr at room temperature with a goat anti-rabbit (1:10,000, Jackson ImmunoResearch Labs, 111035144) or goat anti-mouse (1:5000, Invitrogen, 62-6520) HRP-conjugated secondary antibody in 5% milk in PBST. After three washes with PBST, blots were developed with SuperSignal™ West Pico (Thermo Scientific, 34578) or Femto (Thermo Scientific, 34095) chemiluminescent substrate and imaged on a ChemiDoc Gel Imaging System (Bio-Rad Laboratories). The primary antibodies used in this study are: Mono-ADP-Ribose (AbD33204) (1:250; Bio-Rad Laboratories), GFP (pabg1) (1:2000, Proteintech), HA-Tag (C29F4) (1:1000; Cell Signaling Technologies), RNF114 (GTX107046) (1:1000; Genetex), DTX2 (A7398) (1:500, Abclonal), and RNF166 (AP20484c) (1:1000, Abcepta).

### Sequence and structural analysis

Structural modeling was performed using the AlphaFold3 web interface (Abramson et al., 2024). AlphaFold3 PAE plots were visualized using PAE Viewer (Elfmann & Stulke, 2023). ConSurf analysis of evolutionary conservation was performed using the ConSurf web interface (Ashkenazy et al., 2016; Ashkenazy et al., 2010; Celniker et al., 2013; Glaser et al., 2003; Landau et al., 2005). Structural analysis and figure generation were performed using PyMol (The PyMOL Molecular Graphics System, Version 3.0 Schrödinger, LLC). Sequence alignments were generated using T-Coffee and visualized with JalView (Waterhouse et al., 2009).

## Acknowledgements

The authors wish to thank all members of the Pruneda and Cohen labs for advice and discussion contributing to the experimental design and data analysis for this work. Mass spectrometric analysis was performed within the OHSU Proteomics Shared Resource with partial support from NIH core grants P30EY010572, P30CA069533, and S10RR025571. This work was supported by the National Institute of Neurological Disorders and Stroke (2R01NS088629 to MSC) and by the National Institute of General Medical Sciences (R35GM142486 to JNP).

## Conflict of interest

The authors declare no conflicts of interest.

---

**Supplemental Figure 1.**
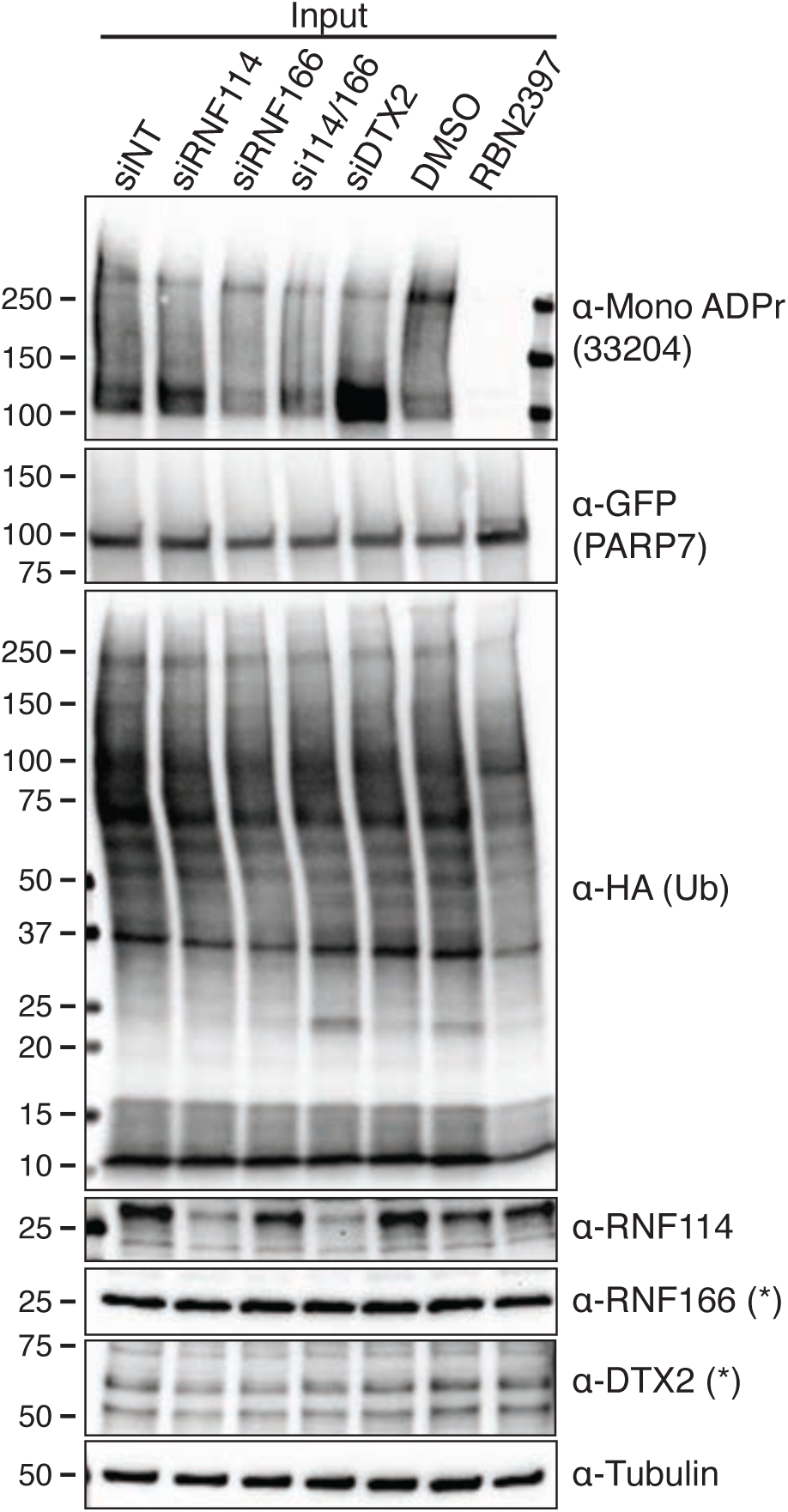
RNF114 extends PARP7 MARUbylation on Glu/Asp residues. Input blots corresponding to Figure 1D prior to the MARUbylation assay to separate PARP7-bound canonical ubiquitylation from MARUbylation. Asterisks indicate blots with high background noise and inability to confirm knockdown of the indicated gene.

**Supplemental Figure 2.**
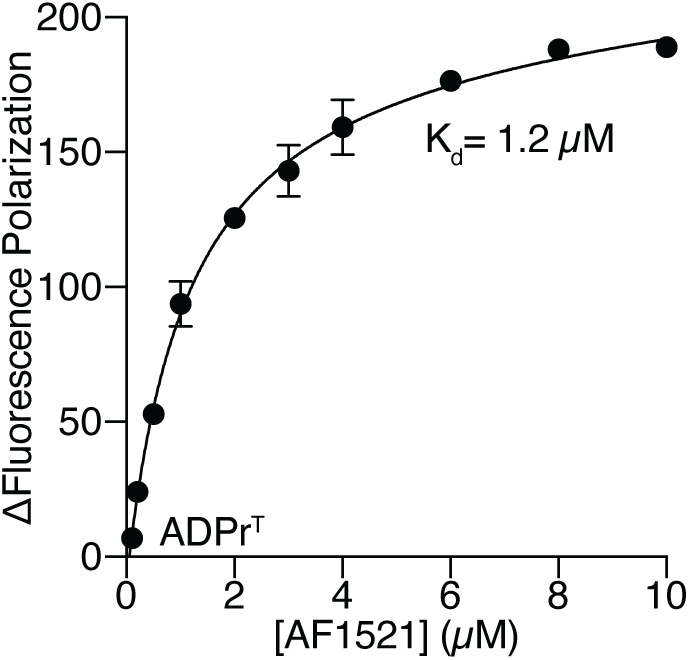
Validation of fluorescent ADPr^T^. Fluorescence polarization binding experiment of ADPr^T^ with the AF1521 macrodomain to validate ADPr^T^ for protein interaction studies.

**Supplemental Figure 3.**
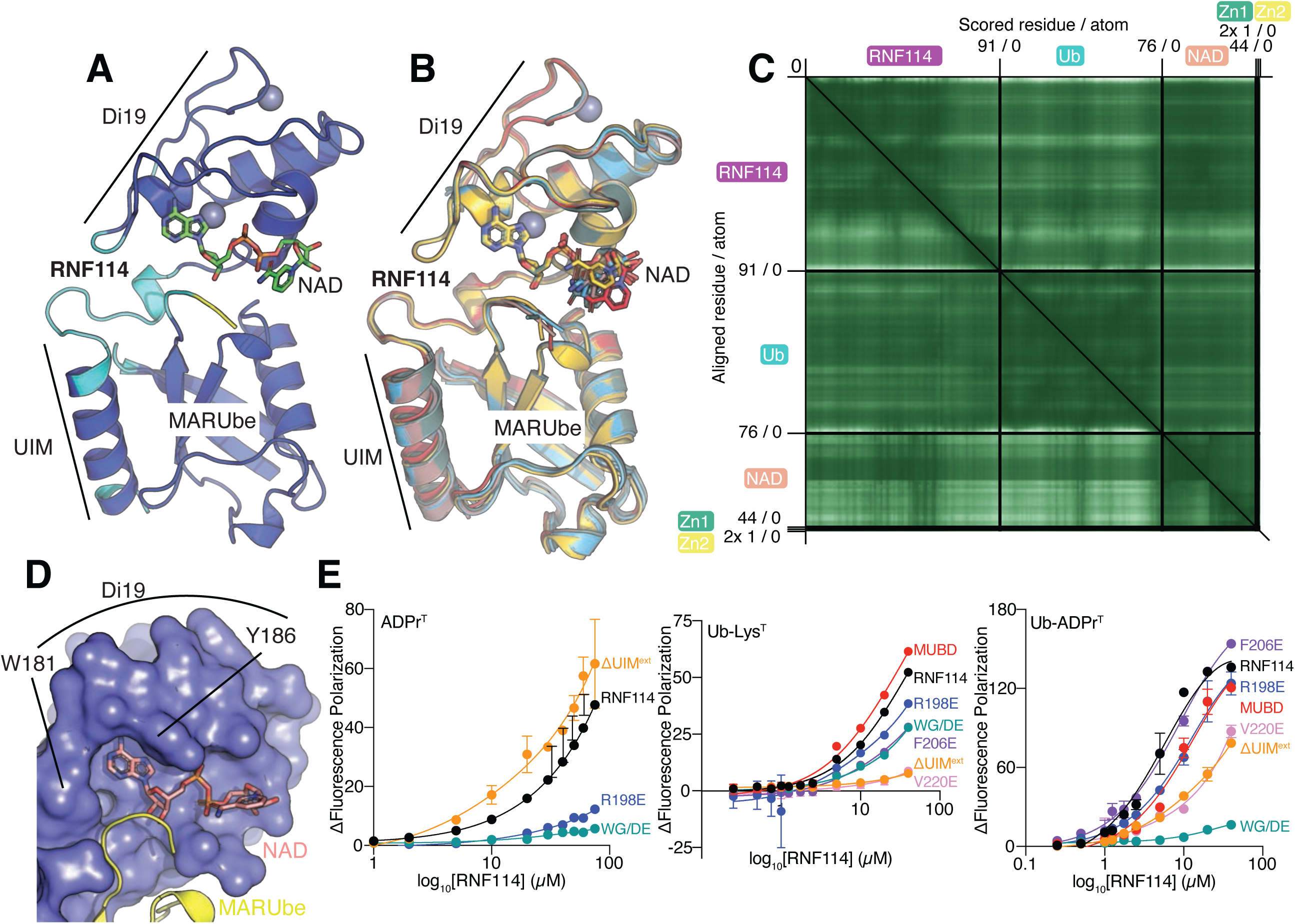
RNF114 binds NAD and Ub. **A)** The MUBD of RNF114 was modeled in AlphaFold3 with two Zn^2+^ ions, NAD, and Ub. The cartoon diagram of the complex is shown with NAD in green and protein colored by AlphaFold3 confidence (pLDDT) where blue represents pLDDT > 90, cyan 70 > pLDDT > 90, yellow 50 > pLDDT > 70, and orange pLDDT < 50. The Di19 domain and UIM that make up the MUBD of RNF114 are labeled. **B**) Superimposition of the top 5 AlphaFold3 models of the MUBD:NAD:Ub complex in different colors. **C**) PAE plot from AlphaFold3 of the MUBD:NAD:Ub model shown in (**A**). Darker green indicates higher confidence in the model. **D**) W181 and Y186 contribute to the positioning of NAD in the binding pocket. Surface representation of the RNF114 Di19 domain accentuates the pocket that accommodates the adenine ring of NAD. **E**) Fluorescence polarization binding curves for the indicated RNF114 mutants with either ADPr^T^, Ub-Lys^T^, or Ub-ADPr^T^ as shown in Figure 3E instead showing the RNF114 concentration (x-axis) on a log scale for ease of comparison.

**Supplemental Figure 4.**
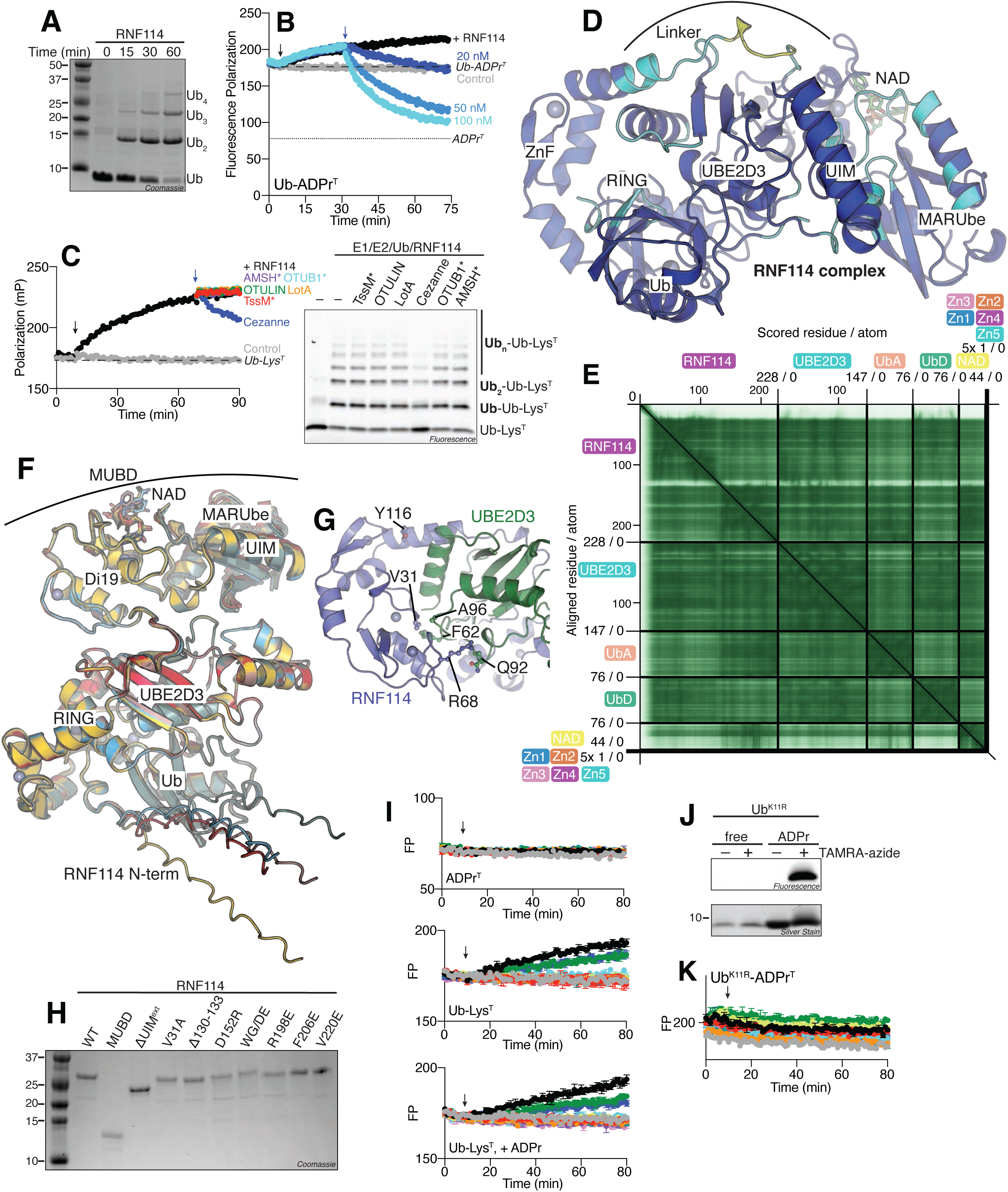
RNF114 catalyzes K11-linked extension on Ub-Lys. **A**) RNF114 autoubiquitylation with WT Ub. Samples were removed at the indicated timepoints and quenched in 3X sample buffer prior to separation by SDS-PAGE and Coomassie staining. **B**) UbiCRest experiment optimizing the concentration of Cezanne to target the K11 isopeptide linkage preferentially over the Ub-ADPr ester linkage. This experiment was conducted as described in the Materials and Methods with the indicated concentrations of Cezanne. **C**) UbiCRest experiment using the indicated deubiquitylases for RNF114 ubiquitylation of Ub-Lys^T^. The reaction was monitored by fluorescence polarization and the sample was removed from the plate at 90 minutes, run on SDS-PAGE, and visualized by in-gel fluorescence of the TAMRA fluorophore. **D**) AlphaFold3 model of the full length RNF114, NAD, UBE2D3, two copies of Ub, and five Zn^2+^ ions. The cartoon diagram of the complex is shown with NAD in green and protein colored by AlphaFold3 confidence (pLDDT) where blue represents pLDDT > 90, cyan 70 > pLDDT > 90, yellow 50 > pLDDT > 70, and orange pLDDT < 50. The RING and ZnF domains and UIM of RNF114, UBE2D3, Ub, the UIM-bound Ub (MARUbe), in addition to the linker of RNF114 that mediates backside UBE2D3 binding are labeled. **E**) PAE plot from AlphaFold3 for the RNF114 transferase complex shown in (**D**). Darker green indicates higher confidence in the model. **F**) The top 5 AlphaFold3 models from (**D**) are shown aligned and colored differently. Each protein and specific regions of RNF114 are labeled. **G**) Detailed view of the modeled interface between the RNF114 RING domain and UBE2D3. V31 and the linchpin residue R68 are shown from the RING domain of RNF114 and their proximity to F62/A96 and Q92, respectively. Y116, a tyrosine phosphorylation site, is also shown in this view. **H**) RNF114 mutants used to evaluate binding and ligase activity were diluted to 1 µM and visualized by SDS-PAGE and Coomassie staining. **I**) UbiReal curves showing the activity of RNF114 mutants against the ADPr^T^ and Ub-Lys^T^ ± unlabeled ADPr substrates. **J**) Validation of the Ub^K11R^-ADPr^T^ substrate (products of the CuAAC reaction) by SDS-PAGE and in-gel fluorescence. **K**) UbiReal experiment for RNF114 mutants against the Ub^K11R^-ADPr^T^ substrate. For all fluorescence polarization panels, the black arrow represents the timepoint when ATP was added to the reaction to initiate RNF114 ligase activity, and the blue arrow represents the timepoint when the indicated deubiquitylase was added to cleave the products.

**Supplemental Figure 5.**
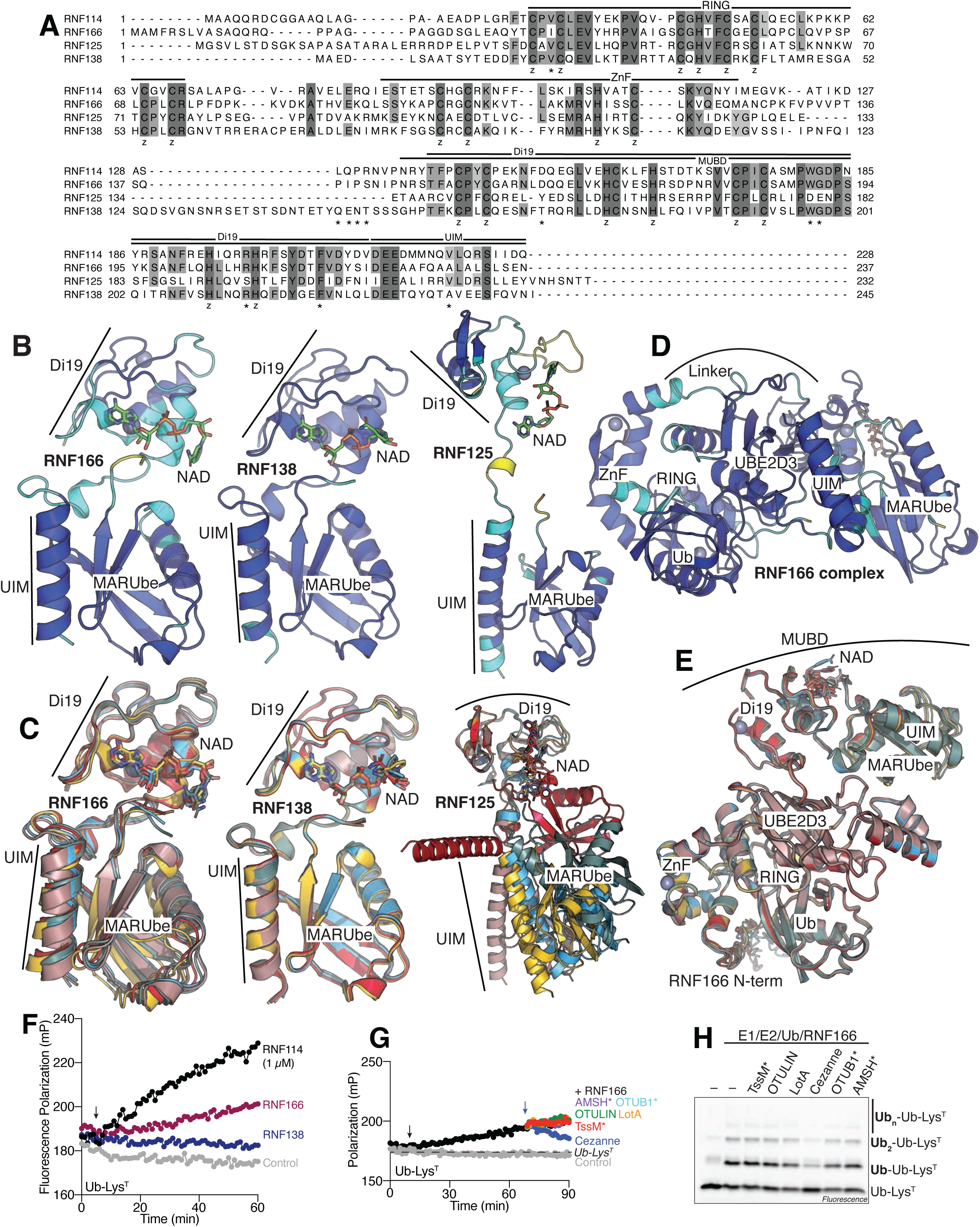
Ub-ADPr recognition is conserved in a family of <u>M</u>AR<u>U</u>be <u>T</u>argeted <u>L</u>igases (MUTLs). **A**) Sequence alignment of MUTLs RNF114, RNF166, RNF125, and RNF138. The RING and ZnF domains, MUBD, Di19, and UIM are labeled based on the sequence of RNF114. Sites of point mutation in our experiments with RNF114 are indicated by asterisks, and Zn^2+^-coordinating residues are labeled (z). Sequences were aligned using Jalview and colored according to percent identity where the darker the grey corresponds to higher conservation. **B**) Cartoon diagrams of the MUBD of RNF166, RNF138, or RNF125 in complex with Ub and NAD colored by the AlphaFold3 confidence scale. **C**) Alignment of the top 5 models for each MUBD (RNF166, RNF138, or RNF125) in complex with NAD and Ub. **D**) AlphaFold3 model of RNF166 in complex with UBE2D3, NAD, two copies of Ub, and five Zn^2+^ ions, colored by confidence. **E**) Overlay of the top 5 AlphaFold3 models for the RNF166 ligase complex. **F**) Representative UbiReal experiment for the indicated E3 ligases with the Ub-Lys^T^ substrate. In this experiment, RNF114 was used at 1 µM and the other ligases were used at 5 µM. ATP addition is marked by the black arrow. **G**) Reaction products of a ligase assay for RNF166 with Ub-Lys^T^ were subjected to a UbiCRest panel. ATP was added to initiate RNF166 ligase activity at the black arrow, and the indicated deubiquitylase was added at the blue arrow. **H**) The 90-minute UbiCRest timepoint from (**G**) was removed from the plate reader and visualized by SDS-PAGE and in-gel fluorescence. (**B, D**) For all AlphaFold3 confidence coloring, blue represents pLDDT > 90, cyan 70 > pLDDT > 90, yellow 50 > pLDDT > 70, and orange pLDDT < 50.

## Notes

### Competing Interest Statement

The authors have declared no competing interest.

