## Supporting Information for "A family of E3 ligases extend K11 polyubiquitin on sites of MARUbylation"

**Method & Materials**

**General chemistry experimentation** 1H NMR were recorded on a Bruker DPX spectrometer at 400 MHz. Chemical shifts are reported as parts per million (ppm) downfield from an internal tetramethylsilane standard or solvent references as s (singlet), d (doublet), t (triplet), q (quartet), p (pentet), h (hextet), hep (heptet), m (multiplet), and br (broad) All reactions were run in flame or oven dried glassware under an atmosphere of dry argon unless otherwise noted Solvents were of ACS chemical grade (Fisher Scientific) and used without further purification unless otherwise indicated. Commercially available starting reagents were used without further purification. Analytical thin-layer chromatography was performed with silica gel 60 F254 glass plates (SiliCycle). Flash column chromatography was conducted with either pre-packed Redisep Rf normal/reverse phase columns Teledyne ISCO) or self-packed columns containing 200-400 mesh silica gel (SiliCycle) on a Combiflash Companion purification system (Teledyne ISCO). High performance liquid chromatography (HPLC) was performed on a Varian Prostar 210 (Agilent) with a flow rate of 20 ml/min using Polaris 5 C18-A columns (150 x 4.6 mm, 3 µm - analytical, 150 x 21.2 mm, 5 µm-preparative) (Agilent). HPLC analytical conditions: mobile phase (MP) A: 50 mM TEAB buffer (aq), mobile phase (MP) B: Acetonitrile; flow rate = 1.0 ml/min; UV-Vis detection: λ1 = 254 nm, λ2 = 220 nm. All final products were ≥95% purity as assessed by this method. Retention times (tR) and purity refer to UV detection at 220 nm. Low-resolution mass spectra were acquired on an Advion Mass-express.

**Scheme 1. Synthesis of O-propargyl RibosePhosphate**

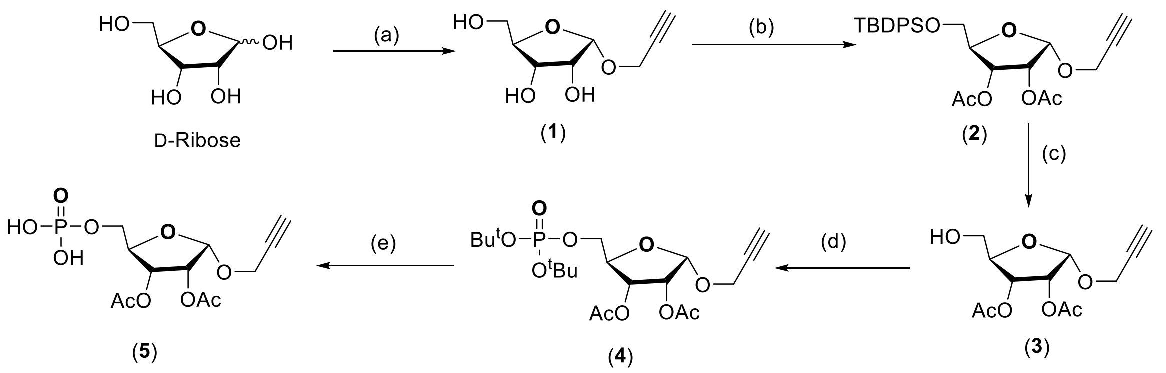


Reagents: (1) Propargyl alcohol, AcCl, Rt; (2) TBDPSCl, Im. DMF, rt; (3) Ac2O, Pyridine, rt; (4) TBAF, THF, rt; (4) Di-*tert*-butyl-*N,N*-diisopropylphosphoramidite, TBHP, ACN; (e) TFA, DCM

**1-O-Propargyl-ribofuranose (1)** [1]

In a 100 ml round bottom flask containing D-Ribose (0.5g, 3.33 mmol) was dissolved in propargyl alcohol (2.5 ml) and cooled down to 0 °C. Addition of acetyl chloride (50 µL) was done in a dropwise manner and then reaction mixture was shifted to room temperature. After stirring at room temperature for 1 hour, the reaction mixture was quenched using pyridine (75 µL) and concentrated using rota-vap. Crude obtained was purified using silica gel flash chromatography using 100% EtOAc as eluent to afford the α (137 mg, 22%) and β (170 mg, 27%) anomers. Spectral data is in agreement as reported in literature.[1]

*α-anomer*: $^{1}$H NMR (400 MHz, MeOD) δ 5.21 (d, *J* = 4.4 Hz, 1H), 4.34 (dd, *J* = 2.5, 1.1 Hz, 2H), 4.08 – 3.99 (m, 2H), 3.96 (dd, *J* = 6.6, 3.3 Hz, 1H), 3.69 (dd, *J* = 12.0, 3.5 Hz, 1H), 3.62 (dd, *J* = 12.0, 4.2 Hz, 1H), 2.85 (t, *J* = 2.4 Hz, 1H).

**α- 1-*O*-Propargyl-2, 3-di-*O*-acetyl-5-*O*-*tert*-butyldiphenylsilyl-ribofuranose (2)**

Compound **1** (0.25g, 1.33 mmol) and Imidazole (0.18g, 2.66 mmol) were dissolved in anhydrous DMF and cooled down to 0 $^{o}$C. Addition of TBDPSCl (0.42 ml, 1.59 mmol) was done in a dropwise manner, after complete addition reaction mixture was shifted to room temperature and stirred for 2hrs. TLC-analysis showed complete consumption of the starting material. Reaction mixture was quenched using saturated $NaHCO_3$ solution (10 ml) and diluted with EtOAc (20ml). Organic layer was collected and dried over Na2SO4, filtered and concentrated under high vacuo to afford crude residue as syrup. The crude obtained was dissolved in anhytdrous pyridine (10 ml) and treated with Acetic anhydride ( 0.5 ml, 5.31 mmol) and catalytic DMAP. Reaction was stirred at room temperature for 16hrs. TLC analysis indicated the complete consumption of the starting material, the reaction mixture was evapotared under high vacuum to afford crude residue. It was diluted with EtOAc and washed with 5% HCl solution, organic layer was collected and dried over $Na_2SO_4$, filtered and concentrated to get crude residue. It was purified using silica gel flash chromatography using 5-20% EtOAc-Hexane as eluent to afford syrup as pure compound as syrup (0.45g, 68%).

$^{1}$H NMR (400 MHz, $CDCl_3$) δ 7.73 – 7.67 (m, 4H), 7.49 – 7.38 (m, 6H), 5.52 (d, *J* = 4.6 Hz, 1H), 5.47 (dd, *J* = 7.0, 2.9 Hz, 1H), 5.21 (dd, *J* = 7.0, 4.6 Hz, 1H), 4.19 (q, *J* = 3.0 Hz, 1H), 4.15 (q, *J* = 7.2 Hz, 1H), 3.90 (dd, *J* = 11.2, 2.9 Hz, 1H), 3.81 (dd, *J* = 11.2, 3.0 Hz, 1H), 2.44 (t, *J* = 2.4 Hz, 1H), 2.17 (s, 3H), 2.14 (s, 3H), 1.09 (s, 9H). ESMS m/z calcd for $C_{28}H_{34}O_7Si$ ([M+H]+ ) 511.21, found 511.21

**1-*O*-propargyl 2,3-di-*O*-acetyl-5-*O*-(di-*tert*-butyl)-phosphoryl-α-ᴅ-ribofuranoside (4)**[2]

Compound **2** (0.44g, 0.86 mmol) was dissolved in anhydrous THF (4.3 ml) and 1M TBAF in THF (1.29 ml, 1.29 mmol) was added. The reaction mixture was stirred at room temperature for 2hrs and then evaporated under rotavap. Crude obtained was diluted with EtOAc (20 ml) and washed with water and brine. Organic layer was collected and dried over $Na_2SO_4$, filtered and concentrated to get crude residue. It was used for the next step without purification.

Compound **3** was co-evaporated with dioxane. To a 50 ml round bottom flask compound **3** (230.0 mg, 0.84 mmol) was taken and a mixture of 1-methyl imidazole.HCl (300 mg, 2.53 mmol) and 1-methyl Imidazole (140 mg, 1.69 mmol) was added. To this reaction mixture di-*tert*-butyl-*N,N*-diisopropylphosphoramidite (0.47 mL, 1.27 mmol) was added and the reaction mixture was stirred at room temperature for 30 min. Then reaction mixture was cooled to 0 °C and then TBHP in decane (0.85 mL, 4.65 mmol, 5.5 M) was added. The solution was allowed to warm to room temperature and stir for 1 h. The reaction was quenched by addition of aq. $NaHCO_3$ and extracted with EtOAc. The organic layer was washed with water and dried over $Na_2SO_4$, concentrated in-vacuo and purified by flash silica gel chromatography using 0-4% DCM/MeOH as eluent to obtain the title compound 4 as a syrup (255 mg, 78% over 2 steps).

$^{1}$H NMR (400 MHz, $CDCl_3$) δ 5.50 (d, *J* = 4.5 Hz, 1H), 5.28 (dd, *J* = 7.2, 3.2 Hz, 1H), 5.06 (dd, *J* = 7.3, 4.5 Hz, 1H), 4.34 (d, *J* = 2.4 Hz, 2H), 4.30 – 4.25 (m, 1H), 4.20 – 4.13 (m, 2H), 2.43 (t, *J* = 2.4 Hz, 1H), 2.14 (s, 6H), 1.51 (s, 18H). $^{13}$C NMR (101 MHz, $CDCl_3$) δ 170.4, 169.7, 98.5, 82.8, 82.8, 82.7, 82.7, 81.1, 81.0, 78.7, 74.70,

70.8, 69.9, 54.5, 29.8, 29.8, 29.8, 29.8, 20.8, 20.5. ESMS m/z calcd for $C_{20}H_{33}O_{10}P$ ([M+H]+ ) 465.19, found 465.19.

Compound **4** (0.24 mg, 0.52 mmol) was dissolved in DCM (5.2 mL) and TFA (0.26 mL) was added. The reaction was stirred at room temperature for 30 minutes, co-evaporated with toluene and pyridine (2x) to afford white solid compound (**5**). The intermediate phosphate was analyzed by mass spectrometry which confirmed the complete removal of the *tert*-butyl group. It was used for the subsequent reaction without further purification.

**Scheme 2: Synthesis of α-*O*-propargyl-ADPr**

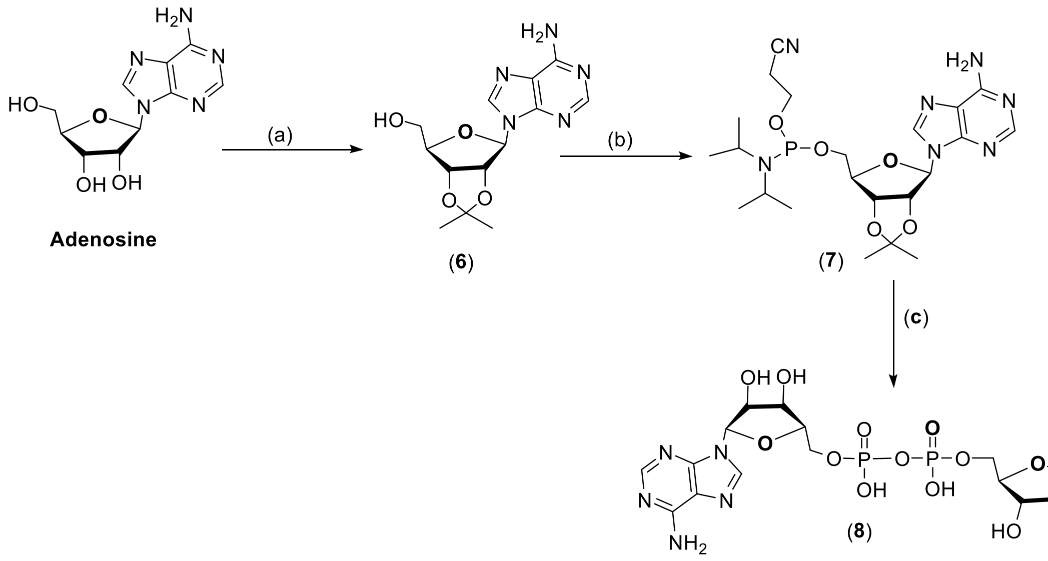


Reagents: (a) *p*TSA, Acetone, rt; (b) 2-cyanoethoxy-*N*,*N*-diisopropylaminochlorophosphine, DIPEA, MeCN; (C) (i) 5, DCI, MeCN, TBHP rt; (ii) DBU; (iii) 50% Formic acid, CAN; (iv) $NH_4OH$, rt

**2', 3'-*O*-Isopropylidene Adensoine (6)**[3]

To a suspension of Adenosine (0.5 g, 1.87 mmol)in dry acetone (100 mL) was added dry TsOH (3.22 g, 18.71 mmol) in two portions. The reaction mixture was stirred under an atmosphere of $N_2$ at room temperature. After 1 hour TLC analysis indicated consumption of the starting material. it was cooled down to 0 °C and quenched with saturated $NaHCO_3$-solution (100 mL) with stirring over 5 minutes. The solvents were removed under reduced pressure and the solid obtained was redissolved in Acetone (100 mL) and stirred overnight. Solid was filtered and filtrate was concentrated to get crude residue. It was purified using silica gel flash chromatography using 0-8% MeOH:DCM as eluent to afford white solid as product (0.41g, 72%). $^1$H NMR (400 MHz, DMSO) δ 8.34 (s, 1H), 8.15 (s, 1H), 7.34 (s, 2H), 6.11 (d, *J* = 3.1 Hz, 1H), 5.34 (dd, *J* = 6.2, 3.1 Hz, 1H), 5.25 (t, *J* = 5.5 Hz, 1H), 4.96 (dd, *J* = 6.2, 2.5 Hz, 1H), 4.21 (td, *J* = 4.9, 2.5 Hz, 1H), 3.61 – 3.46 (m, 2H), 1.54 (s, 3H), 1.32 (s, 3H). ESMS m/z calcd for $C_{13}H_{17}N_5O_4$ ([M+H]+ ) 308.14, found 308.14

**α-*O*-propargyl-ADPr (8)**

To a stirred solution of compound 6 (0.1 g, 0.32 mmol), coevaporated with MeCN, in DCM (2 mL), containing DIPEA (0.14 mL, 0.81 mmol) was added 2-cyanoethoxy-*N*,*N*-diisopropylaminochlorophosphine

(73 µL, 0.325 mmol) under an argon atmosphere. The reaction mixture was stirred for 45 min at room temperature. TLC and mass spectrometry analysis indicated the complete consumption of the starting material and formation of the product. It was evaporated under high vacuum and purged 3 times with argon and used for the next reaction without purification. Intermediate phosphate, 5 (40 mg, 0.11 mmol) and dicyanoimidazole (33.53 mg, 0.28 mmol) were co-evaporated with $CH_3CN$ (3x) and dissolved in dry $CH_3CN$ (1.1 mL). Adenosine amidite 7 (86.5 mg, 0.17 mmol) was added to the reaction mixture, stirred at room temperature for 15 minutes and cooled down to 0 °C. $^tBuOOH$ (5.5 M in decane) (0.11 mL, 0.624 mmol) was added. The reaction was stirred for 30 minutes, TLC analysis indicated the phosphate formation. DBU (91.5 µL, 0.61 mmol) was added and the reaction was stirred for an additional 30 minutes. Solvent was evaporated under rotavap and residue obtained was dissolved using 2:1, CH3CN:H2O (6 mL). 50% formic acid (2mL) was added and the reaction mixture was stirred overnight at room temperature. Solvent was evaporated in-vacuo and co-evaporated twice with $NH_4OH$ to obtain a crude residue. Residue obtained was suspended in CHECN (3 ml) and 35%Ammonium hydroxide (3 mL) was added, the reaction was stirred for 16 hours and concentrated under reduced pressure. The residue obtained was triturated with acetonitrile. The white precipitate was purified by preparative HPLC (Gradient of 100% 50 mM TEAB buffer & 0% to 20% acetonitrile, 50 mM TEAB buffer & 80% acetonitrile). The fractions containing product were pooled and lyophilized to afford pure product 8 (17.1 mg, 21% over 5 steps). Spectral data were found to be in accordance as reported in the literature.[2]

$^1$H NMR (400 MHz, $D_2O$) δ 8.43 (s, 1H), 8.17 (s, 1H), 6.05 (d, *J* = 5.7 Hz, 1H), 5.05 (d, *J* = 3.8 Hz, 1H), 4.69 – 4.65 (m, 1H), 4.44 (dd, *J* = 5.1, 3.7 Hz, 1H), 4.31 (d, *J* = 3.4 Hz, 1H), 4.19 – 4.09 (m, 5H), 4.06 – 4.04 (m, 2H), 3.95 – 3.90 (m, 2H), 2.73 (t, *J* = 2.4 Hz, 1H). $^{13}$C NMR (101 MHz, $D_2O$) δ 155.0, 152.0, 148.9, 140.0, 100.6, 87.0, 83.8, 83.7, 83.6, 79.0, 75.6, 74.3, 70.9, 70.3, 69.4, 65.5, 65.1, 54.8. ESMS m/z calcd for $C_{18}H_{24}N_5O_{14}P_2$ ([M+H]+ ) 598.09, found 598.10

**Scheme 3. Synthesis of TAMRA-ADPr using click reaction**

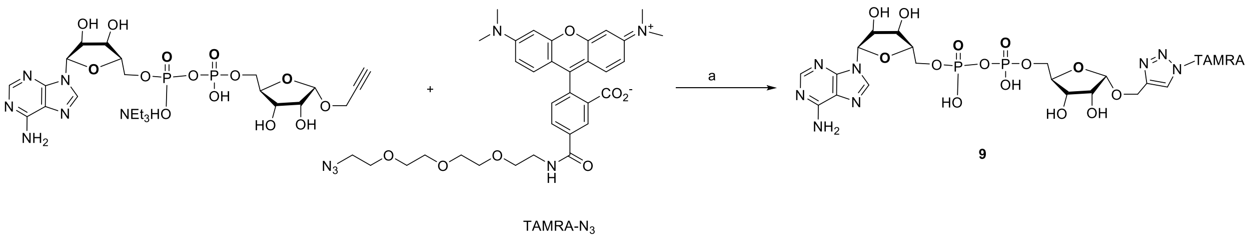


**Reagents used:** (a) $CuSO_4.5H_2O$, NaAsH, $NaHCO_3$, $H_2O$:$^t$BuOH, 1:1

**TAMRA-ADPr Triazole (9)**

To a stirring solution of alkyne (4 mg, 5.7 µmol) in water (0.5 mL) was added $CuSO_4.5H_2O$ (1.1 mg, 4.6 µmol), sodium ascorbate (1.81 mg, 9.2 µmol), $NaHCO_3$ (0.96 mg, 11.5 µmol, 2 eq.) and TAMRA-azide (3.6 mg, 5.7 µmol) in $^t$BuOH (500 µL). The reaction was allowed to proceed at room temperature for overnight before complete conversion of the alkyne was observed by LC-MS. The reaction mixture was concentrated and titurated using CH3CN, centrifuged and solid obtained was dissolved using water. It was subjected to preparative HPLC purification, yielding TAMRA-isoADPr (0.8 mg, 12%) as a dark purple solid. ESMS m/z calcd for $C_{51}H_{63}N_{11}O_{21}P_2$- ([M-H]-) 1226.3, found 1226.3.

**HPLC**

Flow rate: 1.0 mL/min

Solvent A = 50 mM TEAB; Solvent B = Acetonitrile

tR = 9.3 min, 100 % 50 mM TEAB, 0% Acetonitrile to 80% 50 mM TEAB, 20% Acetonitrile over 6 mins; 50% Acetonitrile until 10 mins followed by 50% gradient until 14 mins; 70% Acetonitrile until 14 mins; followed by gradually decreasing to 0% Acetonitrile & 100% 50mM TEAB over 30 mins.

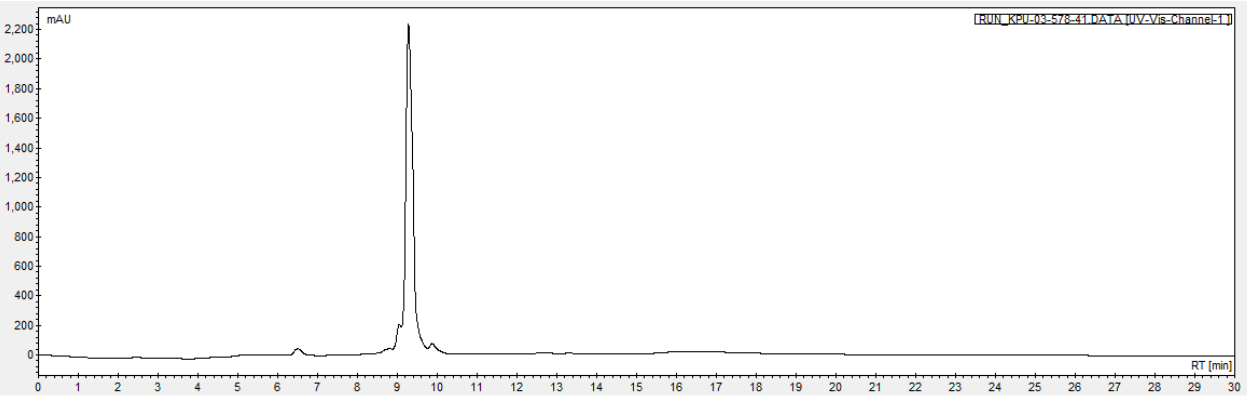

**1H NMR spectrum of α- 1-*O*-Propargyl-ribofuranose (1)**

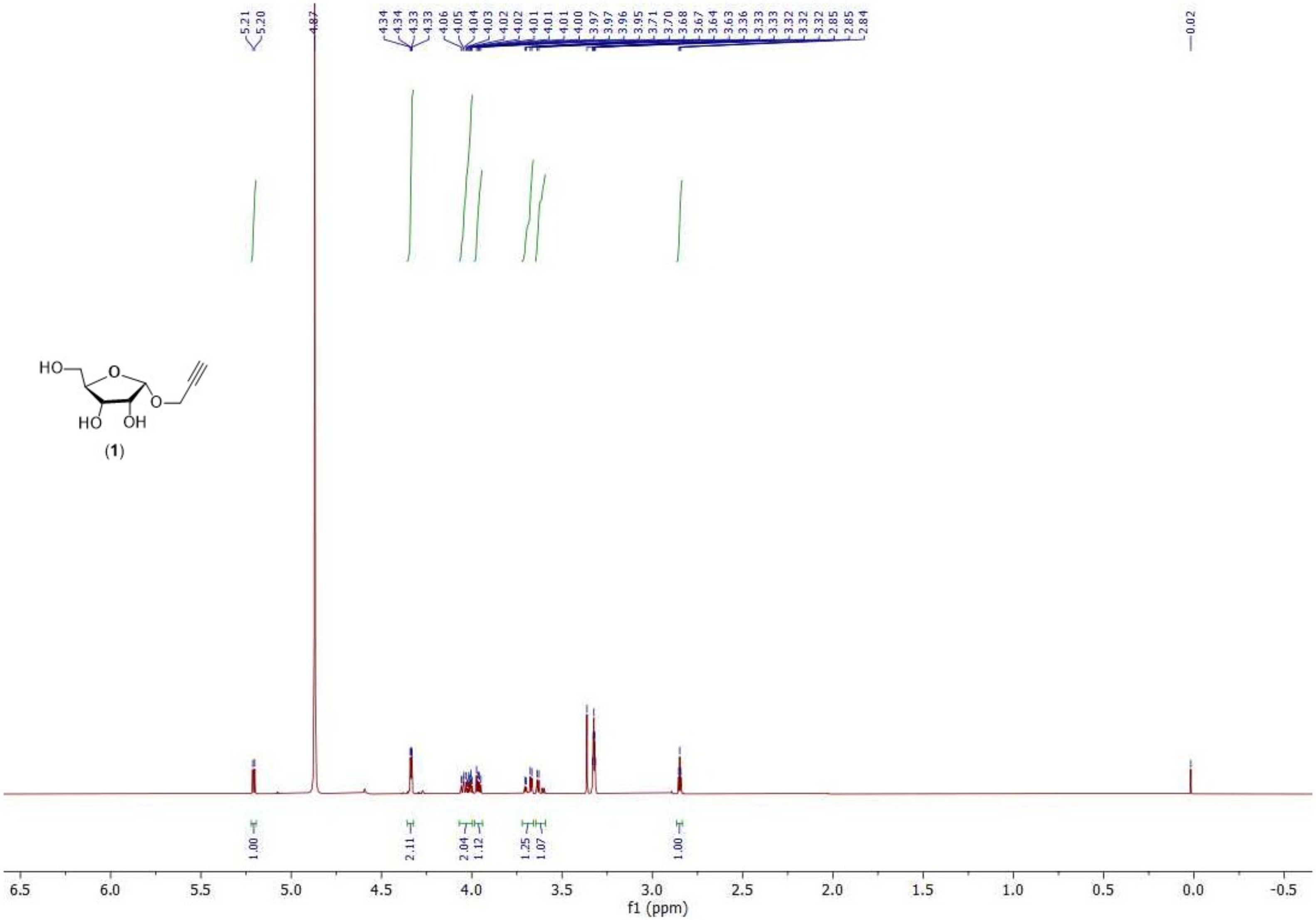

**1H NMR spectrum of α- 1-*O*-Propargyl-2, 3-di-*O*-acetyl-5-*O*-*tert*-butyldiphenylsilyl-ribofuranose (2)**

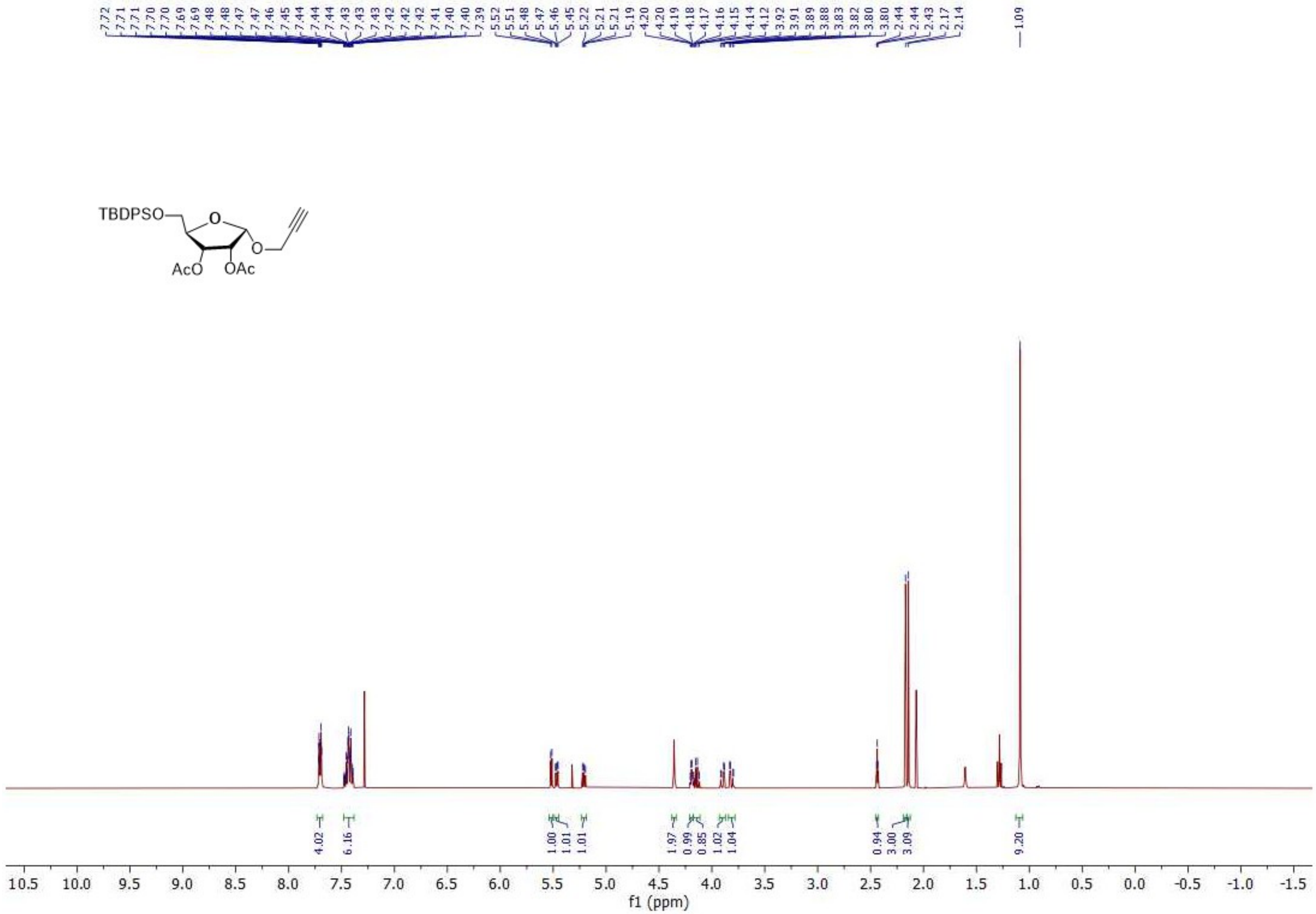

**$^{1}$H NMR spectrum of 1-*O*-propargyl 2,3-di-*O*-acetyl-5-*O*-(di-*tert*-butyl)-phosphoryl-α-ᴅ-ribofuranoside (4)**

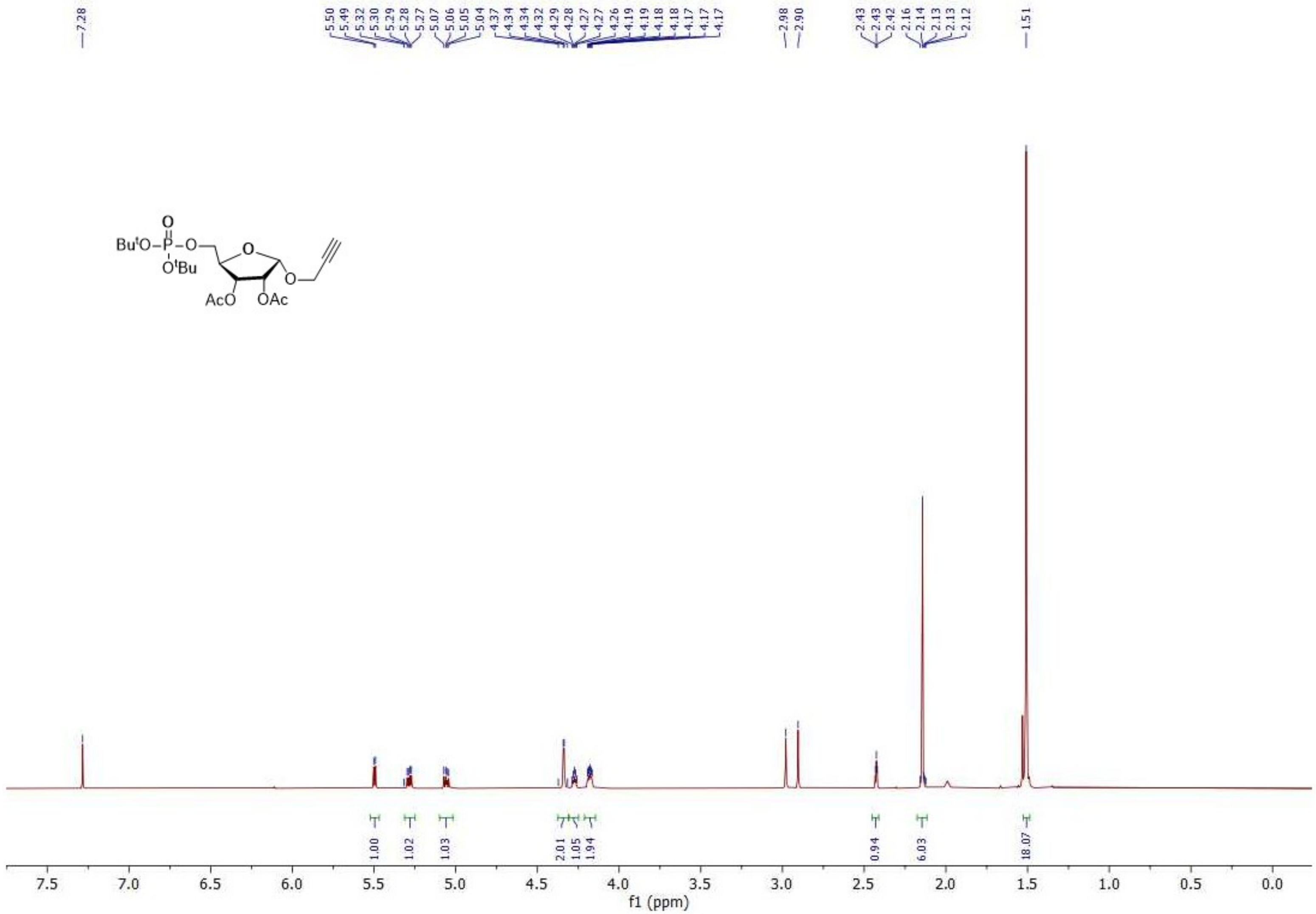

**$^{13}$C NMR spectrum of 1-*O*-propargyl 2,3-di-*O*-acetyl-5-*O*-(di-*tert*-butyl)-phosphoryl-α-ᴅ-ribofuranoside (4)**

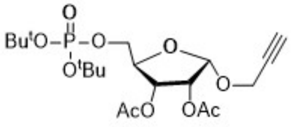


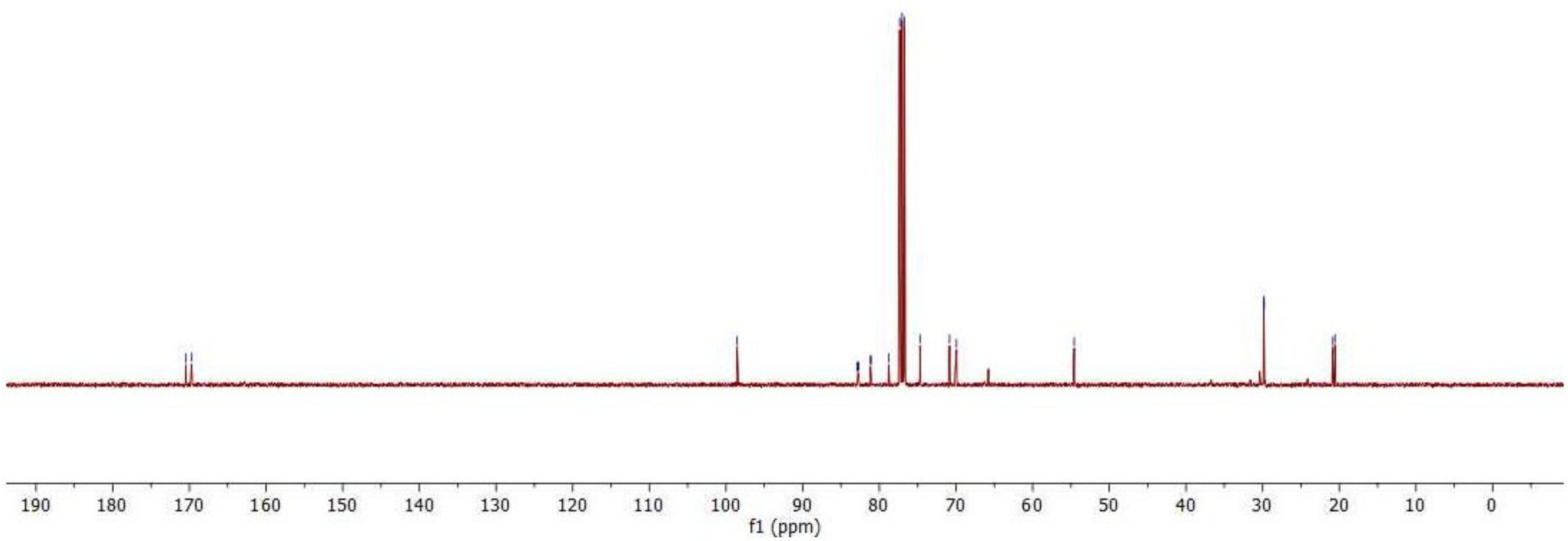

**$^{1}$H NMR spectrum 2′, 3′-*O*-Isopropylidene Adenosine (6)**

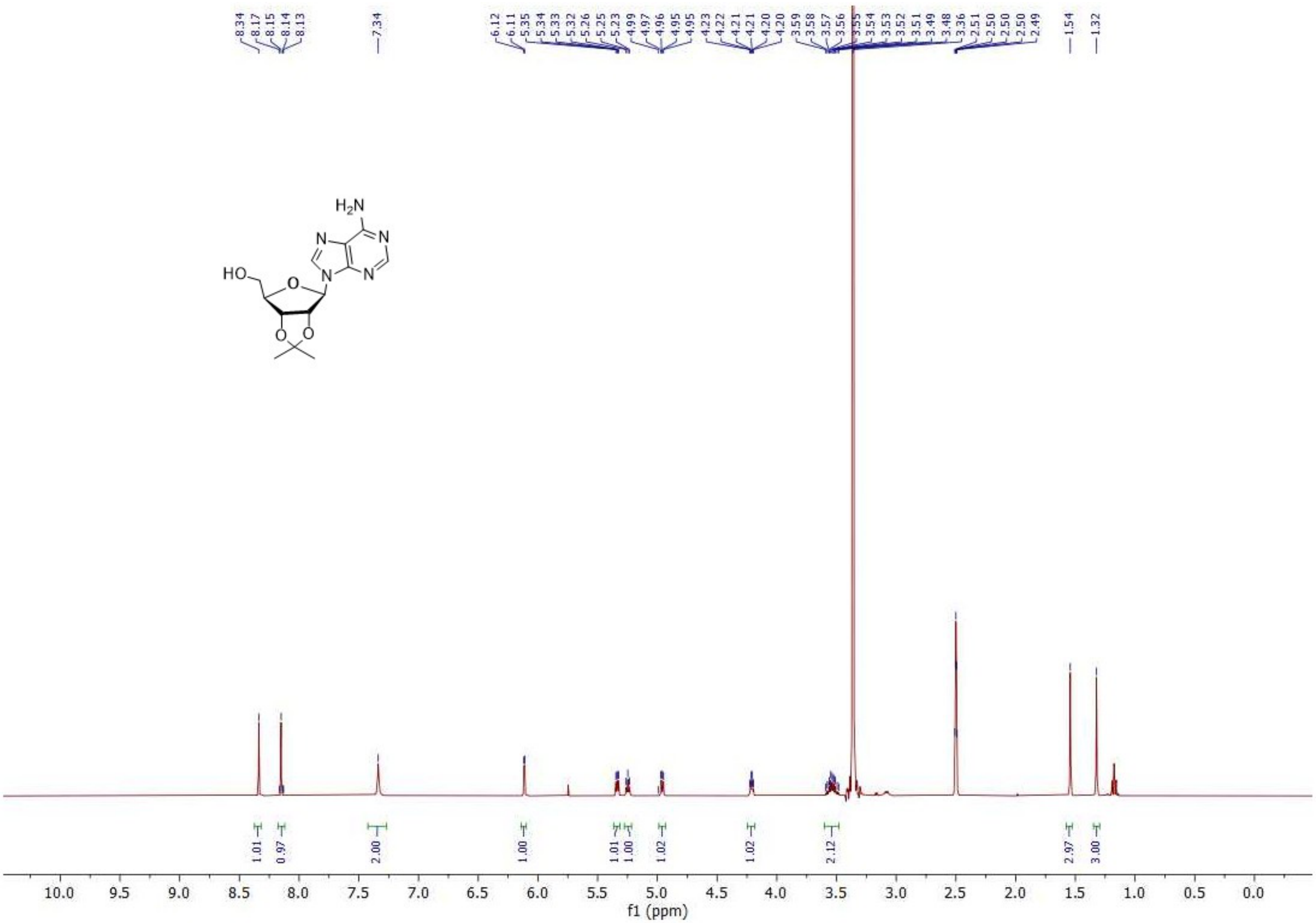

**1H NMR spectrum of α-*O*-propargyl-ADPr (8)**

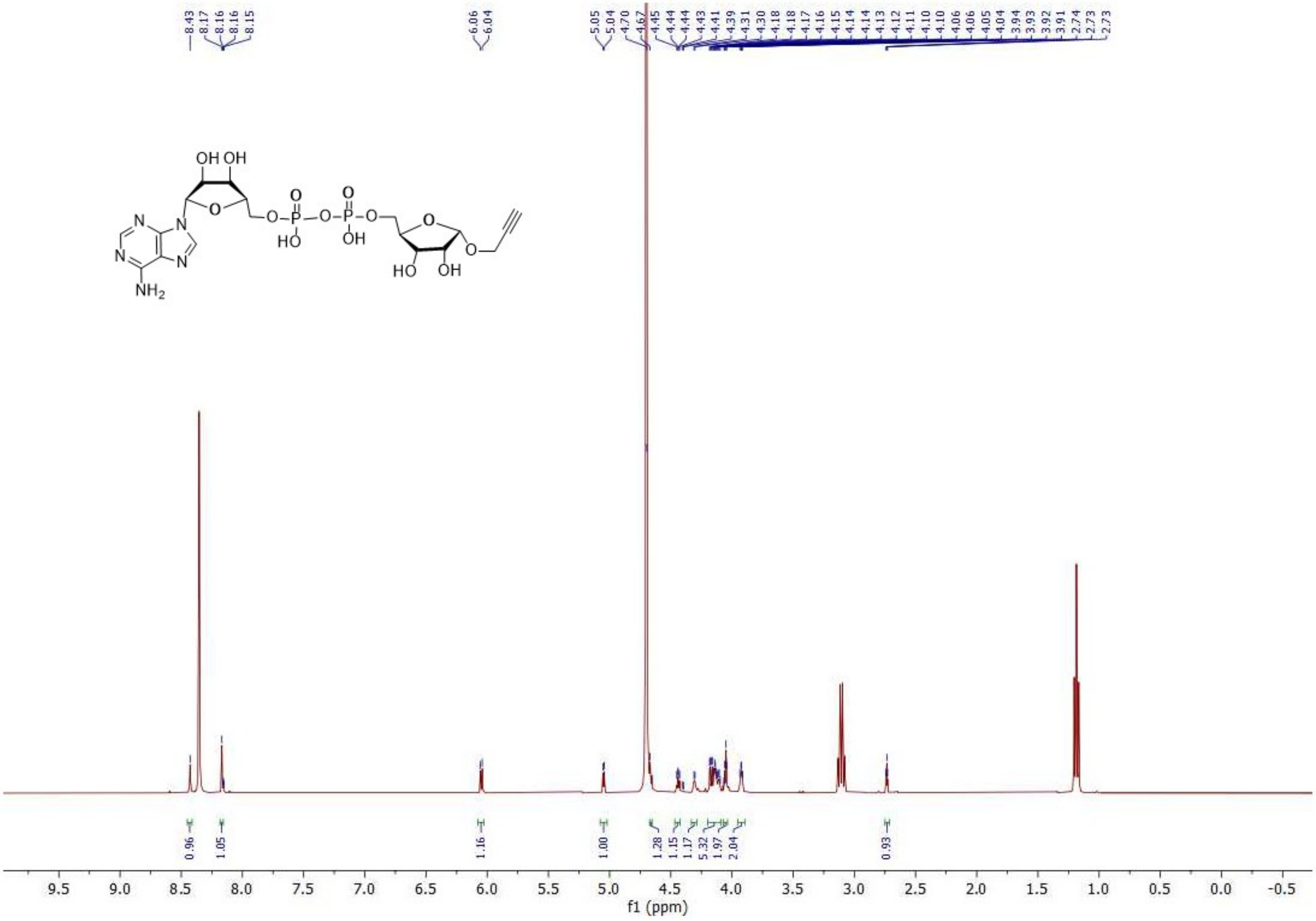

**13C NMR spectrum of α-*O*-propargyl-ADPr (8)**

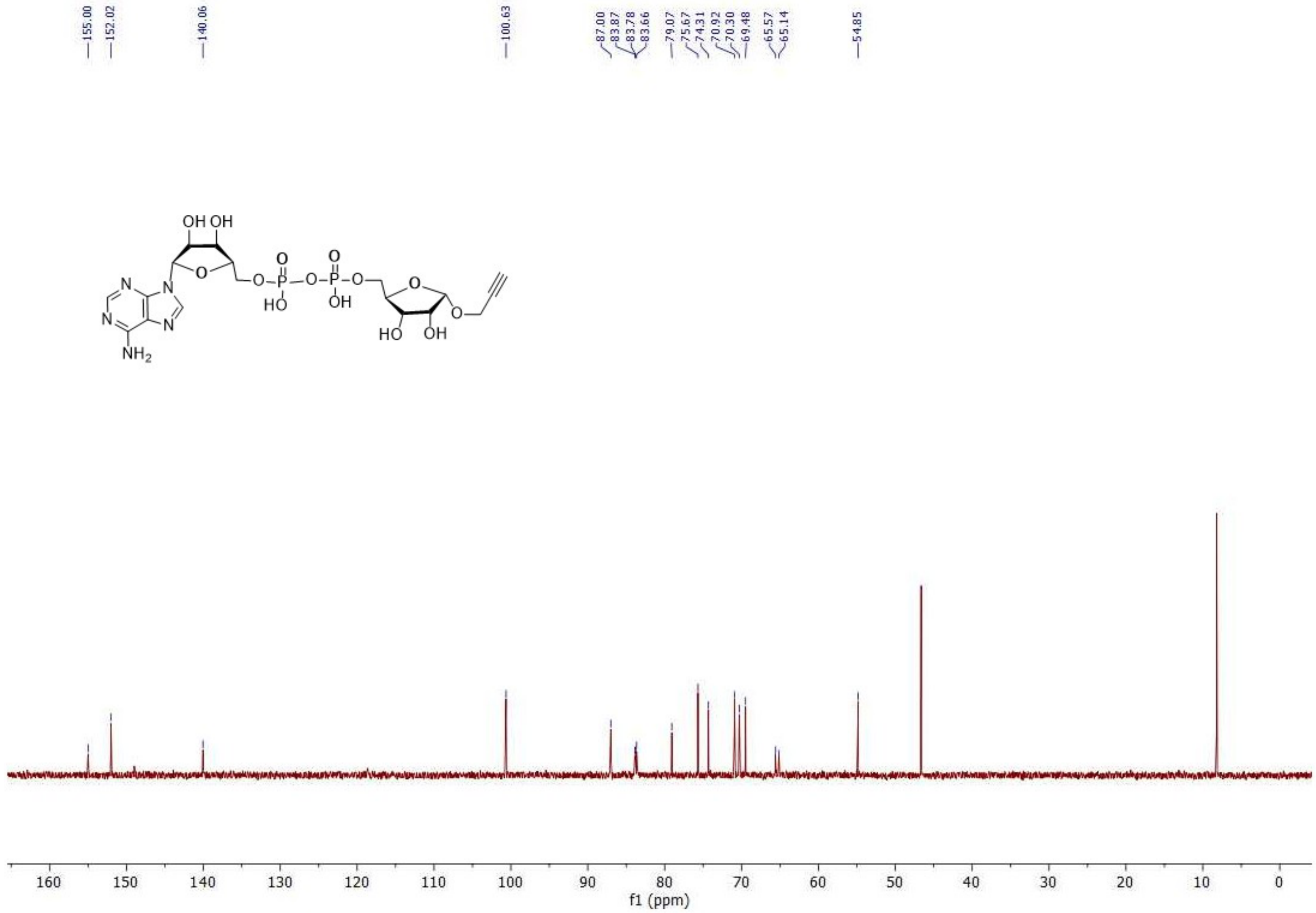

**COSY NMR spectrum of α-*O*-propargyl-ADPr (8)**

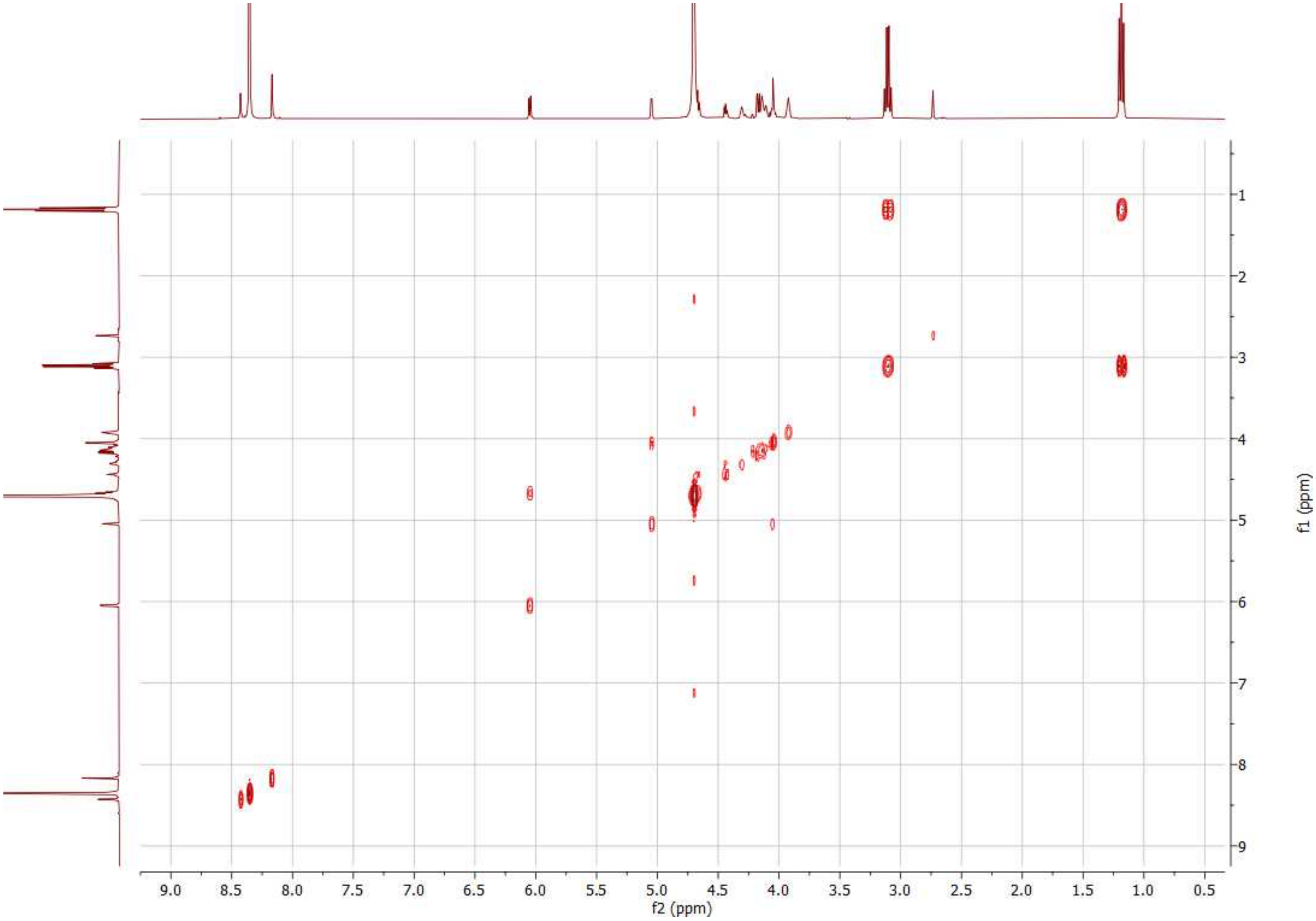

**HSQC NMR spectrum of α-*O*-propargyl-ADPr (8)**

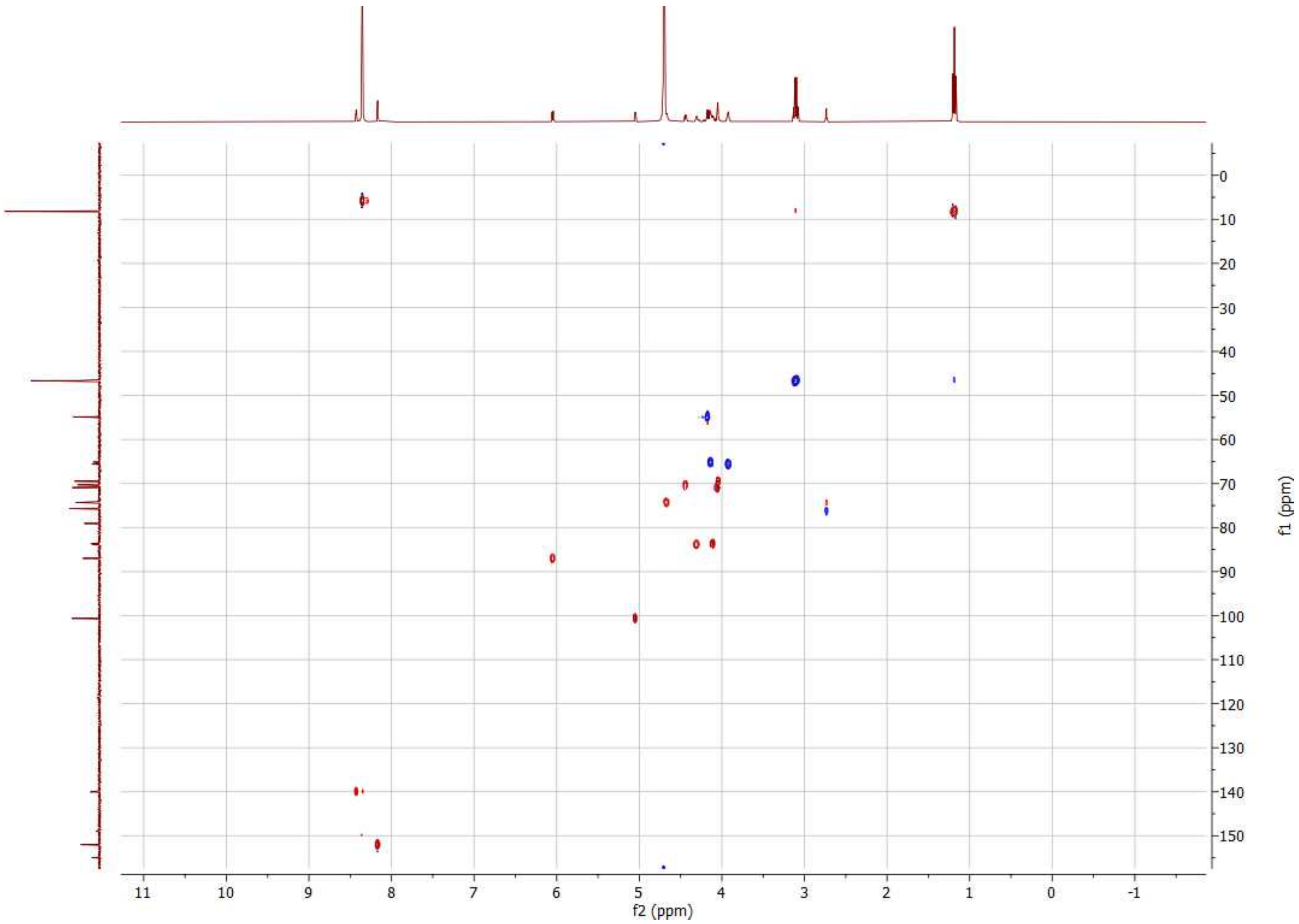
